# A proposed new *Tombusviridae* genus featuring extremely long 5’ untranslated regions and a luteo/polerovirus-like gene block

**DOI:** 10.1101/2024.06.23.600130

**Authors:** Zachary Lozier, Lilyahna Hill, Elizabeth Semmann, W. Allen Miller

## Abstract

*Tombusviridae* is a large family of single-stranded, positive-sense RNA plant viruses with uncapped, non-polyadenylated genomes encoding 5-7 open reading frames (ORFs). Previously, we discovered, by high-throughput sequencing of maize and teosinte RNA, a novel genome of a virus we call Maize-associated tombusvirus (MaTV). Here we determined the precise termini of the MaTV genome by using 5’ and 3’ rapid amplification of cDNA ends (RACE). In GenBank, we discovered eleven other nearly complete viral genomes with MaTV-like genome organizations and related RNA-dependent RNA polymerase (RdRp) sequences. These genomes came from diverse plant, fungal, invertebrate and vertebrate organisms, and some have been found in multiple organisms across the globe. The available 5’ untranslated regions (UTRs) of these genomes are remarkably long: at least 438 to 727 nucleotides (nt), in contrast to those of other tombusvirids, which are <150 nt. Moreover these UTRs contain 6 to 12 AUG triplets that are unlikely to be start codons, because - with the possible exception of MaTV - there are no large or conserved ORFs in the 5’ UTRs. Such features suggest an internal ribosome entry site (IRES), but we found no conserved secondary structures. In the 50 nt upstream of and adjacent to the ORF1 start codon, the 5’ UTR was cytosine-rich and guanosine-poor. As in most tombusvirids, ORF2 (RdRp gene) appears to be translated by in-frame ribosomal readthrough of the ORF1 stop codon. Indeed, in all twelve genomes we identified RNA structures known in other tombusviruses to facilitate this readthrough. ORF5 is predicted to be translated by readthrough of the ORF3 (coat protein gene) stop codon as in genus *Luteovirus*. The resulting readthrough domains are highly divergent. ORF4 overlaps with ORF3 and may initiate with a non-AUG start codon. We also found no obvious 3’ cap-independent translation elements, which are present in other tombusvirids. The twelve genomes diverge sufficiently from other tombusvirids to warrant classification in a new genus. Because they contain two leaky stop codons and a potential leaky start codon, we propose to name this genus *Rimosavirus* (*rimosa* = leaky in Latin).

## 1 Introduction

Metagenomics has revolutionized virus discovery. Searching for viruses by sequencing total RNA from environmental samples (metagenomics), such as soil (Nicolas et al., 2023), seawater (Culley et al., 2006), or organisms has resulted in an exponential increase in known viruses, or viral genomes in the past decade (Roossinck, 2017;Harvey and Holmes, 2022). The viruses associated with these newly discovered genomes are mostly uncharacterized, but the obvious viral nature of the genomes indicate that those viruses exist (Simmonds et al., 2017). Because virus particles can be highly abundant and stable, viruses isolated from an organism may not actually infect that organism, but may be just “hitching a ride”. For example, plant viruses have been identified in human intestinal microbiome (Tiamani et al., 2022), bat guano (Li et al., 2010), and in aphid vectors that transmit them (Feng et al., 2017) but which are not infected by them. Hence, viruses known only by sequence and the organism from which they are isolated are labeled as “associated” with that organism.

In large scale metagenomics projects, thousands of viral genome sequences have been automatically assembled, annotated, and deposited in GenBank, in some cases with very little direct scrutiny by humans (Shi et al., 2016;Yang et al., 2022). Because of the numerous noncanonical translation mechanisms used by many RNA viruses, these autoannotated genomes are often mis-annotated (Cobbin et al., 2021). Also, if the viral genomes are not of interest to the sequencers, or if the sequencers simply lack the time to study these viral genomes, they may remain essentially undiscovered in GenBank. Here we describe several genomes of viruses in the *Tombusviridae* family (tombusvirids) that appear to fall in this category.

The *Tombusviridae* family contains over 100 virus species (tombusvirids) in eighteen genera officially named by the International Committee on Virus Taxonomy (ICTV). This large and diverse family includes many economically costly pathogens, such as maize chlorotic mottle virus, which has devastated maize production in mixed infection with a potyvirus in East Africa (Redinbaugh and Stewart, 2018), and the barley yellow dwarf viruses which comprise the most ubiquitous viruses of wheat, barley and oat, worldwide (Trebicki et al., 2015;Peters et al., 2022). In addition, tombusvirids have proved to be excellent models. For example, the first X-ray crystal structure of an icosahedral virus was that of tomato bushy stunt virus (for which the family is named) (Winkler et al., 1977). Also the roles of host proteins and subcellular structures in every stage of the virus life cycle are better understood for TBSV than almost any other RNA virus (Nagy, 2016;Chkuaseli and White, 2018;Nagy and Feng, 2021). Tombusvirids contain a positive strand RNA genome of 4-6 kb which lacks a 5’ cap and a poly(A) tail, as the genome terminates in CCC (White and Nagy, 2004;Simon, 2015). *Dianthovirus* is the only tombusvirid genus that has a bipartite genome (Okuno and Hiruki, 2013).

The tombusvirid genome encodes 5-7 open reading frames (ORFs). ORF1 encodes a replication protein that lines membrane-bound replication vesicles (Nagy and Feng, 2021). ORF2 encodes the RNA-dependent RNA polymerase (RdRp) and is translated as a long C-terminal extension of ORF1 via ribosomal readthrough of – or frameshift around – the ORF1 stop codon (Barry and Miller, 2002;Cimino et al., 2011;Tajima et al., 2011;Kuhlmann et al., 2016). Downstream ORFs encode movement proteins, suppressors of RNA silencing, coat protein and vector transmission components (Chay et al., 1996;Kong et al., 1997;Lakatos et al., 2004;White and Nagy, 2004). They are translated - often via various leaky start and stop codons (Dinesh-Kumar and Miller, 1993;Johnston and Rochon, 1996;Cimino et al., 2011;Chkuaseli and White, 2022) - from subgenomic mRNAs that are 5’-truncated versions of genomic RNA (Wang and Simon, 1997;Koev and Miller, 2000;Jiwan and White, 2011). To allow translation of the uncapped genomic and subgenomic mRNAs, a 3’ cap-independent translation element (3’ CITE) is present in the 5’ end of the 3’ UTR (Fabian and White, 2004;Simon and Miller, 2013). The 3’ CITE is followed at the 3’ end by RNA structures and sequences required for RNA replication (Simon, 2015), and in some cases structures that regulate readthrough or frameshifting at the ORF1 stop codon via long-distance base pairing (Miller and White, 2006;Nicholson and White, 2014).

After discovery by deep sequencing of the genome of the novel tombusvirid maize-associated tombusvirus (MaTV) in maize and teosinte leaves (Lappe et al., 2022), we searched GenBank for related sequences. Here we describe eleven other viral genomes, all found by metagenomic sequencing, that are related to the MaTV. Based on (i) genome organization, (ii) sequences of RNA-dependent RNA polymerase (RdRp) and coat protein (CP), and (iii) conserved RNA secondary structures, all of these viruses clearly belong in the *Tombusviridae* family.

However their sequences and genome features, such as extremely long 5’ UTR, place them in a clade sufficiently distinct to merit classification in a new genus.

## 2 Materials and Methods

### 2.1 Rapid Amplification of cDNA Ends (RACE)

Sequences of oligonucleotides used for RACE are listed in Supplementary Table S1.

*Source of RNA.* The RNA used for RACE was from the same total RNA extract from maize leaf that was used for sequencing the MaTV genome (Lappe et al., 2022). The leaf was collected from an unhealthy, possibly diseased maize plant near Irapuato, Mexico in October 2017. Total RNA was extracted using a Zymo Direct-zol RNA Miniprep Plus kit (Zymo Research, Irvine, CA, USA), depleted for ribosomal RNA using an Illumina Ribo-Zero rRNA Removal Kit (Plant Leaf) (Illumina, San Diego, CA, USA), concentrated with a Zymo RNA Clean & Concentrator-5 kit, and stored at -80C. Details are described in Lappe et al. (2022).

*First-strand cDNA synthesis for 5’ RACE.* Using Millipore Sigma’s 5’/3’ RACE Kit, 2^nd^ Generation, MaTV cDNA was synthesized using random primers on 1 µg of the above total maize leaf RNA. Following the kit’s instructions, the reaction mixture was incubated at 55 ^O^C for 60 minutes and for another 5 minutes at 85 ^O^C. Immediately after first-strand cDNA synthesis, the cDNA products were purified using Sigma’s High Pure PCR Product Purification Kit, following the specific instructions for cDNA purification outlined in the 5’ RACE protocol as opposed to the protocol that comes with the purification kit.

*PCR on the cDNA sample for 5’ RACE.* PCR was carried out following the purification step after first-strand cDNA synthesis, using Sigma-Aldrich’s Expand High Fidelity PCR System. Using the Expand High Fidelity Buffer with 15 mM MgCl2, thermal cycler PCR conditions were set according to Sigma’s RACE protocol with an altered annealing temperature to accommodate both the MaTV-specific primer and the oligo dT anchor primer in the RACE kit. Three rounds of PCR were conducted on the samples using a different nested primer each round (Nested primers 1, 2, and 3), without purifying the PCR product or diluting the PCR product before the next round was started. The final PCR product was subjected to gel electrophoresis and gel extraction followed by ethanol precipitation and resuspension in 50 µl of nuclease-free water. The resulting DNA was sequenced at the Iowa State University DNA Facility using MaTV-specific 5’ RACE sequencing primer, 5R1 (Supplementary Table S1).

*Ligation of an artificial poly(A) tail to MaTV RNA for 3’ RACE.* The Millipore Sigma 3’ RACE kit utilizes a poly(A) tail on the RNA sample. Because viruses in the family *Tombusviridae*, which includes MaTV, do not have a poly(A) tail, an oligonucleotide (3’poly(A)) containing a poly(A) tract was ligated onto the 3’ end of the total RNA sample using NEB’s T4 RNA Ligase 1 kit. This sequence is complementary to the oligo d(T) anchor primer from the RACE kit and was designed with both a 5’ and 3’ phosphate to prevent the oligo from ligating to itself and ligating to RNA in the total RNA sample that already possesses a poly(A) tail. Using 1 µl of the total RNA sample including Thermo Fisher’s RNaseOut RNA inhibitor, the reaction mixture was incubated at 25 ^O^C for 2 hours and the reaction was stopped by column cleanup using the NEB Monarch RNA Cleanup Kit.

*First-strand cDNA synthesis for 3’ RACE.* Following Millipore Sigma’s 5’/3’ RACE Kit, 2^nd^ Generation, 5ul of the newly polyadenylated total RNA sample containing the ligated oligo(dT) anchor primer was used at a concentration of 100 ng/µl. After incubation at 55 ^O^C for 60 min immediately followed by incubation at 85 ^O^C for 5 min, the cDNA was ready for PCR. No purification step was necessary after cDNA synthesis.

*PCR on the cDNA sample for 3’ RACE.* PCR was carried out immediately following first-strand cDNA synthesis using Sigma-Aldrich’s Expand High Fidelity PCR System. Using the Expand High Fidelity Buffer with 15 mM MgCl2, thermal cycler conditions were set according to Sigma’s RACE protocol altered to accommodate both the MaTV-specific primer, 3F1 (Supplementary Table S1), and the oligo dT anchor primer in the RACE kit. The PCR product was ethanol precipitated and resuspended in 50 µl of nuclease-free water.

*Gel electrophoresis and gel extraction.* 16 µl of purified PCR product at a concentration of 100 ng/µl was run on a 2% agarose gel for 30 minutes at 150V to isolate the MaTV cDNA produced from the total RNA sample. A visible band at ∼400 bp (estimated product size at the 3’ end of MaTV genome) was gel extracted using Qiagen’s QIAquick PCR & Gel Cleanup Kit. After measuring the concentration of cDNA in the eluted sample after gel extraction (∼5 ng/µl), an additional round of PCR was conducted to increase the cDNA concentration (∼300 ng/µl).

*3’ end sequencing of MaTV.* Following PCR and purification of the MaTV cDNA, the samples were sent to Iowa State University’s DNA Sequencing Facility for dideoxy sequencing using MaTV- specific primer 3F2 (Supplementary Table S1) that was nested 3’ of the MaTV-specific primer used for PCR.

### 2.2 Multiple sequence alignment and phylogeny prediction

The RNA-dependent RNA polymerase (RdRp), coat protein (CP), and read-through domain (RTD) amino acid sequences of select viruses from families *Tombusviridae* and Solemoviridae were aligned with Muscle v3.8.31 using default parameters in SnapGene. RNA alignments were imaged using Jalview (Waterhouse et al., 2009;Procter et al., 2021). For phylogenetic tree construction, multiple sequence alignments were passed to FastTree 2.1.11 with the flags “-lg -gamma” to predict phylogenies where branch split reliability values are estimated with the Shimodaira-Hasegawa test (Shimodaira and Hasegawa, 1999). The resultant trees were drawn with FigTree v1.4.4 and re-rooted using the nearest relative outside of the taxonomic group of the compared sequences (DeSalle et al., 2023): Providence virus (NC_014126.1) for the RdRp tree and Ourmia melon virus (NC_011070.1) for the coat protein tree. Accession numbers are the GenBank accession numbers from which the respective amino acid sequences were taken.

### 2.3 RNA structures

RNA secondary structures were predicted using MFOLD (Zuker, 2003), Vienna package (Lorenz et al., 2011), Scanfold2 (Andrews et al., 2018) under default parameters, in iterations with multiple sequence alignments and inspection with the human eye. Secondary structures were drawn using RNAcanvas (Johnson and Simon, 2023).

## 3 Results

### 3.1 Identification of viruses similar to MaTV: a new genus

Previously, we discovered and assembled genomic sequences of MaTV from maize (GenBank no. OK0180181) and teosinte (OK0180182). These genomes are 99.5% identical and none of the base differences affect lengths of open reading frames (ORFs) (Lappe et al., 2022). Thus they are isolates of the same virus; for all subsequent comparisons in this paper, we use the maize isolate. To identify other viruses related to MaTV, we performed a BLAST search of GenBank seeking matches with the RNA-dependent RNA polymerase (RdRp) gene (ORF2) of MaTV. We use the RdRp because a major component of RNA virus classification is based on sequence similarities of the RdRps, as it is the key replication protein encoded by all RNA viruses (Dolja et al., 2020;Simmonds et al., 2023). We also searched for genes with similarity to MaTV ORF4 which seemed to be unique to this genus, when compared to known tombusvirids.

The BLAST searches revealed nearly complete genomes of eleven other viruses with substantial sequence similarity in the RdRp ORF and similar genome organizations to that of MaTV (Fig. 1). This includes apple virus E (AVE), which we reported previously as being similar to MaTV (Lappe et al., 2022). Like the MaTV genome, all eleven of these genomes were discovered in metagenomics sequencing projects using Illumina sequencing (Shi et al., 2016;Chiapello et al., 2020;Waller et al., 2022;Yang et al., 2022;Ni et al., 2023). As discussed below, all of the genomes were misannotated because ORFs were assumed to begin with an AUG codon. Yet, by comparison with other members of the *Tombusviridae*, there appear to be two ORFs (ORFs 2 and 5) that are translated via in-frame readthrough of the stop codon of the preceding ORF. Thus, these ORFs begin immediately after the stop codon rather than at the first AUG codon of the ORF. Because these genomes were found in large, exploratory metagenomics sequencing projects, we do not know what hosts they infect or if any symptoms were associated with these viruses, but they have been found associated with a remarkable variety of organisms including plants, fungi, invertebrates and vertebrates (Table 1). Some virus species have been found more than once, in organisms of diverse kingdoms across the globe (Table 1).

**Fig. 1.**
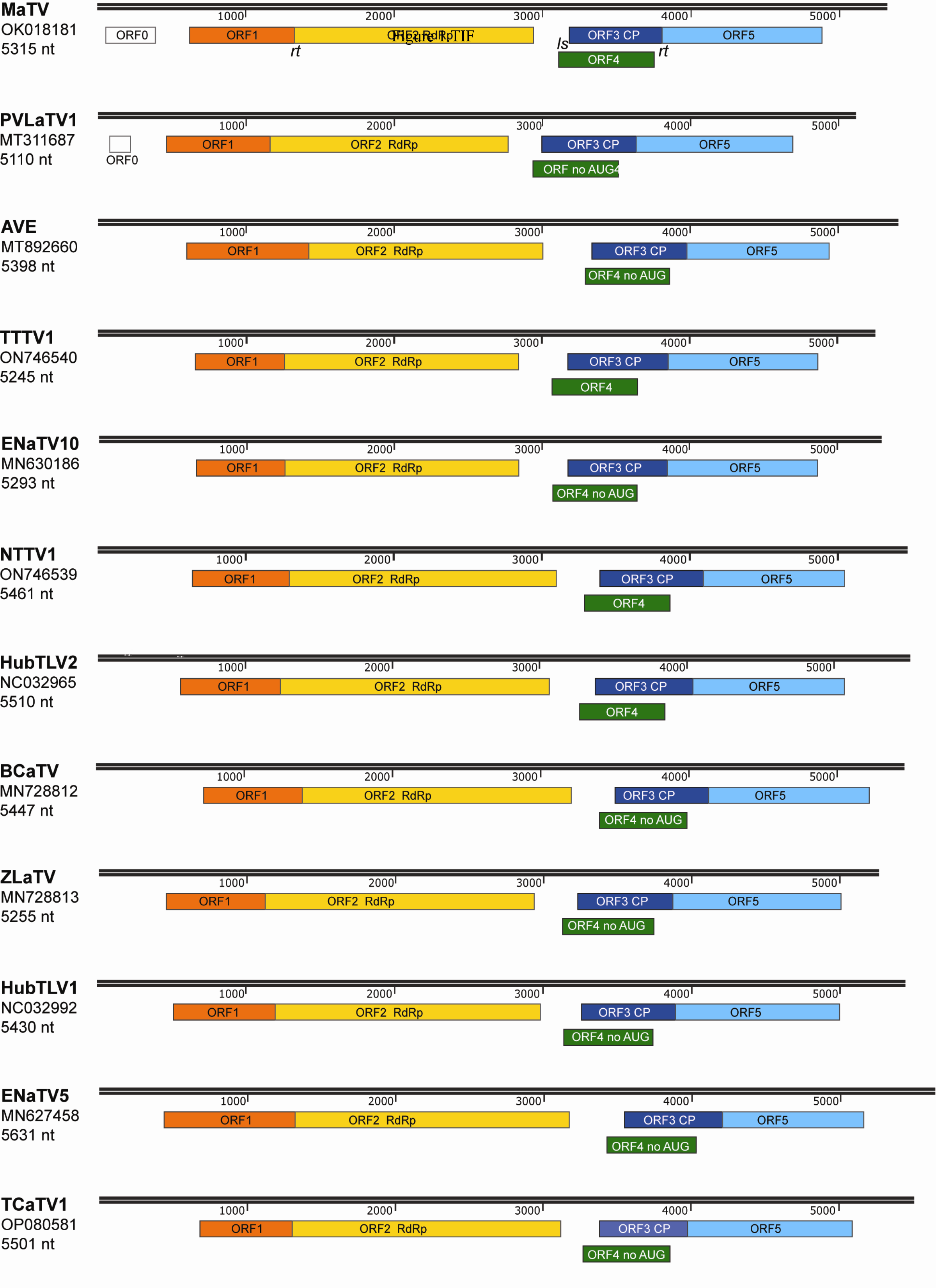
Maps of genomes in proposed new genus, *Rimosavirus.* Virus acronym, GenBank accession numbers used throughout this manuscript, and length of genome sequence for that accession number are shown at left. Colored boxes indicate ORFs, with functions of the protein products of ORF2 (RdRp) and ORF3 (CP) indicated. Scale bar (in nt) is indicated in black above each genome map. ORFs 2 and 5 are translated by in-frame readthrough (rt) of the ORF1 and ORF3 stop codons, respectively. ORFs 4 of the indicated viruses (no AUG) lack an in-frame AUG start codon upstream of ORF3 AUG start codon (see text), which is predicted to be translated froma a subgenomic mRNA via leaky scanning (ls).

**Table 1.**
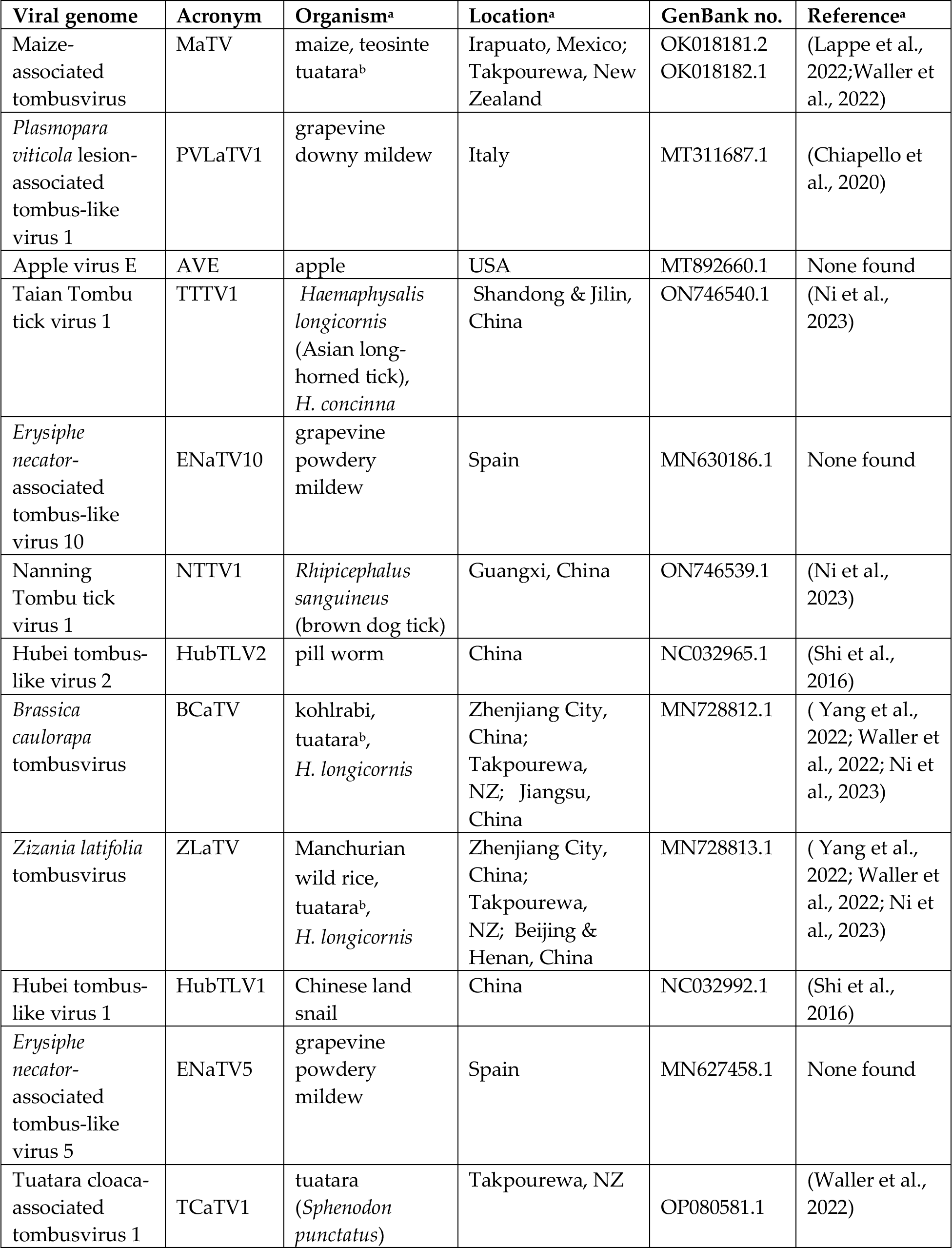
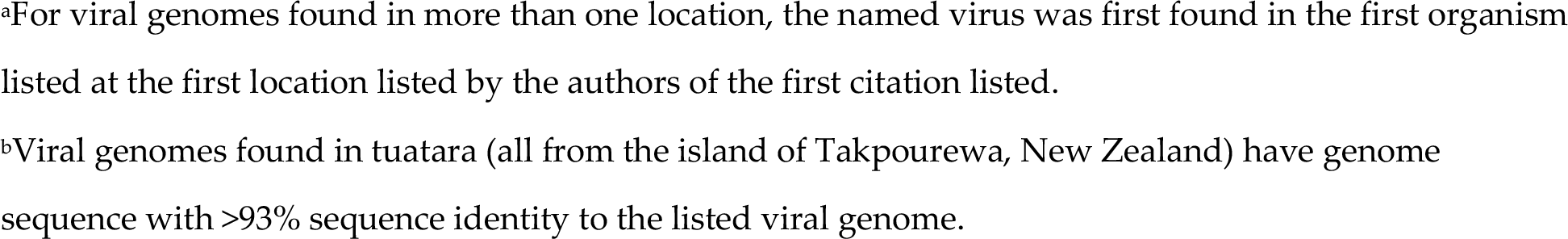
Genomes and sources of viruses in this study.

| **Viral genome** | **Acronym** | **Organism<sup>a</sup>** | **Location<sup>a</sup>** | **GenBank no.** | **Reference<sup>a</sup>** |
| --- | --- | --- | --- | --- | --- |
| Maize-associated tombusvirus | MaTV | maize, teosinte tuatara<sup>b</sup> | Irapuato, Mexico; Takpourewa, New Zealand | OK018181.2<br>OK018182.1 | (Lappe et al., 2022; Waller et al., 2022) |
| *Plasmopara viticola* lesion-associated tombus-like virus 1 | PVLaTV1 | grapevine downy mildew | Italy | MT311687.1 | (Chiapello et al., 2020) |
| Apple virus E | AVE | apple | USA | MT892660.1 | None found |
| Taian Tombu tick virus 1 | TTTV1 | *Haemaphysalis longicornis* (Asian long-horned tick), *H. concinna* | Shandong & Jilin, China | ON746540.1 | (Ni et al., 2023) |
| *Erysiphe necator*-associated tombus-like virus 10 | ENaTV10 | grapevine powdery mildew | Spain | MN630186.1 | None found |
| Nanning Tombu tick virus 1 | NTTV1 | *Rhipicephalus sanguineus* (brown dog tick) | Guangxi, China | ON746539.1 | (Ni et al., 2023) |
| Hubei tombus-like virus 2 | HubTLV2 | pill worm | China | NC032965.1 | (Shi et al., 2016) |
| *Brassica caulorapa* tombusvirus | BCaTV | kohlrabi, tuatara<sup>b</sup>, *H. longicornis* | Zhenjiang City, China; Takpourewa, NZ; Jiangsu, China | MN728812.1 | ( Yang et al., 2022; Waller et al., 2022; Ni et al., 2023) |
| *Zizania latifolia* tombusvirus | ZLaTV | Manchurian wild rice, tuatara<sup>b</sup>, *H. longicornis* | Zhenjiang City, China; Takpourewa, NZ; Beijing & Henan, China | MN728813.1 | ( Yang et al., 2022; Waller et al., 2022; Ni et al., 2023) |
| Hubei tombus-like virus 1 | HubTLV1 | Chinese land snail | China | NC032992.1 | (Shi et al., 2016) |
| *Erysiphe necator*-associated tombus-like virus 5 | ENaTV5 | grapevine powdery mildew | Spain | MN627458.1 | None found |
| Tuatara cloaca-associated tombusvirus 1 | TCaTV1 | tuatara (*Sphenodon punctatus*) | Takpourewa, NZ | OP080581.1 | (Waller et al., 2022) |

<sup>a</sup>For viral genomes found in more than one location, the named virus was first found in the first organism listed at the first location listed by the authors of the first citation listed.
<sup>b</sup>Viral genomes found in tuatara (all from the island of Takpourewa, New Zealand) have genome sequence with >93% sequence identity to the listed viral genome.

In addition to high sequence similarity in the RdRp ORF, these genomes share the following features. (i) They contain an extremely long putative 5’ untranslated region (UTR) ranging from at least 438 nt to 727 nt, although most are almost certainly longer than listed because it is unlikely that most have been sequenced to the 5’ end (below); (ii) The presumed first translated ORF (ORF1, MW 23-34 kDa, Supplementary Table S2) is followed immediately by ORF2, which encodes the RdRp (59-69 kDa) in an arrangement that suggests translation of ORF2 via ribosomal readthrough of the ORF1 stop codon. This arrangement is present in 16 of the 18 *Tombusviridae* genera, the exceptions being *Luteovirus* and *Dianthovirus* genera, members of which employ a -1 ribosomal frameshift for translation of the RdRp. (iii) ORF2 is followed by a noncoding region followed by ORF3, which encodes the coat protein (CP, 22-26 kDa), and an overlapping ORF4 (∼17-20 kDa) (Fig. 1, Supplementary Table S2). The stop codon of ORF3 is followed immediately in the same reading frame by ORF5, suggesting translation of ORF5 via readthrough of the ORF3 stop codon to generate a 34-41 kDa C-terminal extension to the CP (Fig. 1, Supplementary Table S2). This arrangement of ORFs 3, 4, and 5 resembles that of viruses in the genera *Luteovirus* (*Tombusviridae*) and *Polerovirus* (*Solemoviridae*) (Miller and Lozier, 2022), with the exception that ORF4 likely initiates upstream of ORF3 in these new viruses. In all luteo- polero- and enamoviruses (L/P/E viruses), ORF5 has been shown to be translated via readthrough of the ORF3 stop codon (Brown et al., 1996;Xu et al., 2018b;Chkuaseli and White, 2022), and the protein product of ORF5 (the readthrough domain – RTD) is required for aphid transmission (Brault et al., 1995;Chay et al., 1996;Brault et al., 2005;Schiltz et al., 2022), and participates in virus cell to cell movement in the luteo- and poleroviruses (Mutterer et al., 1999;Brault et al., 2000;Peter et al., 2008). In the new tombusvirids reported here, ORF5 is followed by a 3’ UTR with minimum lengths of 229 to 458 nt, but it is unlikely that the 3’ UTR sequences are complete all the way to the 3’ end, except for those of MaTV and possibly *Erysiphe necator*-associated tombus-like virus 10 (ENaTV10) (discussed below).

Whole genome comparisons revealed that Taian tombu tick virus 1 (TTTV1) and ENaTV10 are 95% identical, thus they are strains of the same virus even though one was obtained from an invertebrate parasite of mammals and the other from a plant pathogenic fungus. MaTV and *Plasmopara viticola* lesion-associated tombus-like virus 1 (PVLaTV1) genomes also show close similarity at 68.5% nucleotide sequence identity (73.4% identity in the RdRp ORF), but are clearly different species based on ICTV demarcation criteria for tombusvirids, which is <85% amino acid (aa) sequence identity in the CP, as the CPs show 71.2% aa identity (76.6% nt identity).

We propose that these twelve viruses comprise a new genus in the *Tombusviridae* family. This is based on the distance of the clade in the phylogenetic tree of RdRp sequences from the nearest relative, oat necrotic dwarf virus (genus *Avenavirus*) (Fig. 2), and their distinctive genome organizations, which differ in the same ways from those of other tombusvirids. Because all twelve of these viruses have two probable leaky stop codons (at the ends of ORFs 1 and 3), and may initiate translation of the ORF3 via ribosomal leaky scanning, we propose to call this new genus *Rimosavirus* (*rimosa,* Latin for leaky). While this name is only provisional, for convenience throughout this manuscript, we refer to the twelve viruses on which this paper focuses (Table 1) as rimosaviruses.

**Fig. 2.**
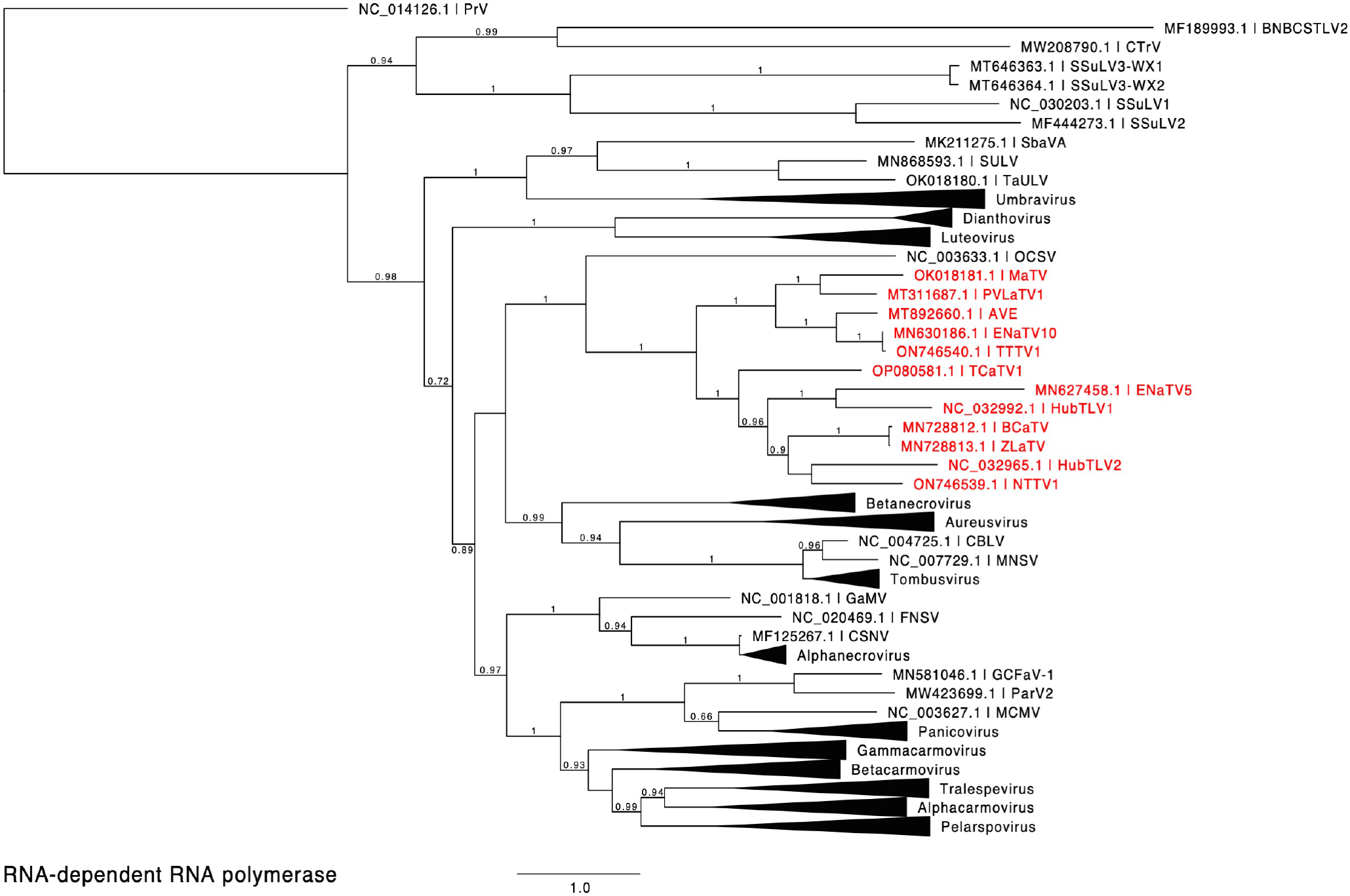
Phylogenetic tree predicting the relationship of viruses based on the amino acid sequences of full-length RdRps (ORF1-ORF2 fusion products). Red entries indicate those sequences belonging to the proposed new *Rimosavirus* genus. Branch support values are shown for splits > 0.5 and are calculated from 1,000 resamples of the Shimodaira-Hasegawa test (SH-like local supports). Branch lengths indicate arbitrary units of evolutionary distances. Providence virus (PrV) was used as outgroup because it is the nearest relative outside of the *Tombusviridae* (DeSalle et al., 2023). For individual viruses (single member genus or unassigned to genus), GenBank accession numbers and virus acronyms are shown.

### 3.2 5’ and 3’ untranslated regions

As mentioned above, it is not clear if any of the rimosaviral genomes were sequenced completely to the 5’ and 3’ ends. To determine the terminal sequences of MaTV, we performed 5’ and 3’ RACE (rapid amplification of cDNA ends)(Yeku and Frohman, 2011) on the same preparation of total maize RNA from which the nearly complete MaTV genome sequence was obtained. RACE revealed 19 additional nt at the 5’ end (GAAAAUAUUUAGGGUACUA) and 62 nt at the 3’ end (UUCCAAACUGCUCAGUAAUGAGAACUUCAAUUACAGUACAGCUAGACAGAUCUGUAAUGCCC) that were not found in the Illumina sequence assembly (accession no. OK018181.1) (Lappe et al., 2022). That sequence has been updated to include the above RACE results (accession no. OK018181.2). The complete genome length is 5315 nt with 622 nt upstream of ORF1, which we call the 5’ UTR (below), and a 434 nt 3’ UTR. Assuming the same length of sequence is present at the ends of the 99% identical teosinte isolate, the GenBank sequence (OK018182.1) lacks 20 nt and 50 nt at the 5’ and 3’ ends, respectively, but we have not performed 5’ or 3’ RACE on that sample.

#### 3.2.1 5’ untranslated region

The 5’ end of the MaTV genome starts with GAAAAUAU, which is highly similar to that of apple virus E (AVE), GAAAAUCU. These sequences resemble the 5’ ends of other tombusvirids which start with a purine (usually a G) followed by a purine-rich, C-poor tract. Thus, the GenBank sequence of AVE (accession no. MT892660) appears to have a complete 5’ end. The GenBank sequence of PLVaTV1 (MT311687) appears to be missing 18 nt from its 5’ terminus, based on alignment with its close relative MaTV (Supplementary Fig. S1). The 5’ ends of the other nine rimosavirus sequences found in GenBank do not begin with a similar sequence, and alignments (Supplementary Fig. S1) suggest numerous bases are missing from the 5’ ends.

The 5’ termini of characterized tombusvirid genomes form a stem-loop of modest stability (Guo et al., 2001;Chattopadhyay et al., 2011). We predicted the secondary structures of the 5’- terminal 40 nt of MaTV and AVE genomes and indeed found such stem-loops (Fig. 3A). For comparison, the known terminal structures of tombusvirids barley yellow dwarf virus (BYDV, *Luteovirus*) and saguaro cactus virus (SCV, *Carmovirus*) are shown. This further supports that the MaTV and AVE 5’ ends are complete.

**Fig. 3.**
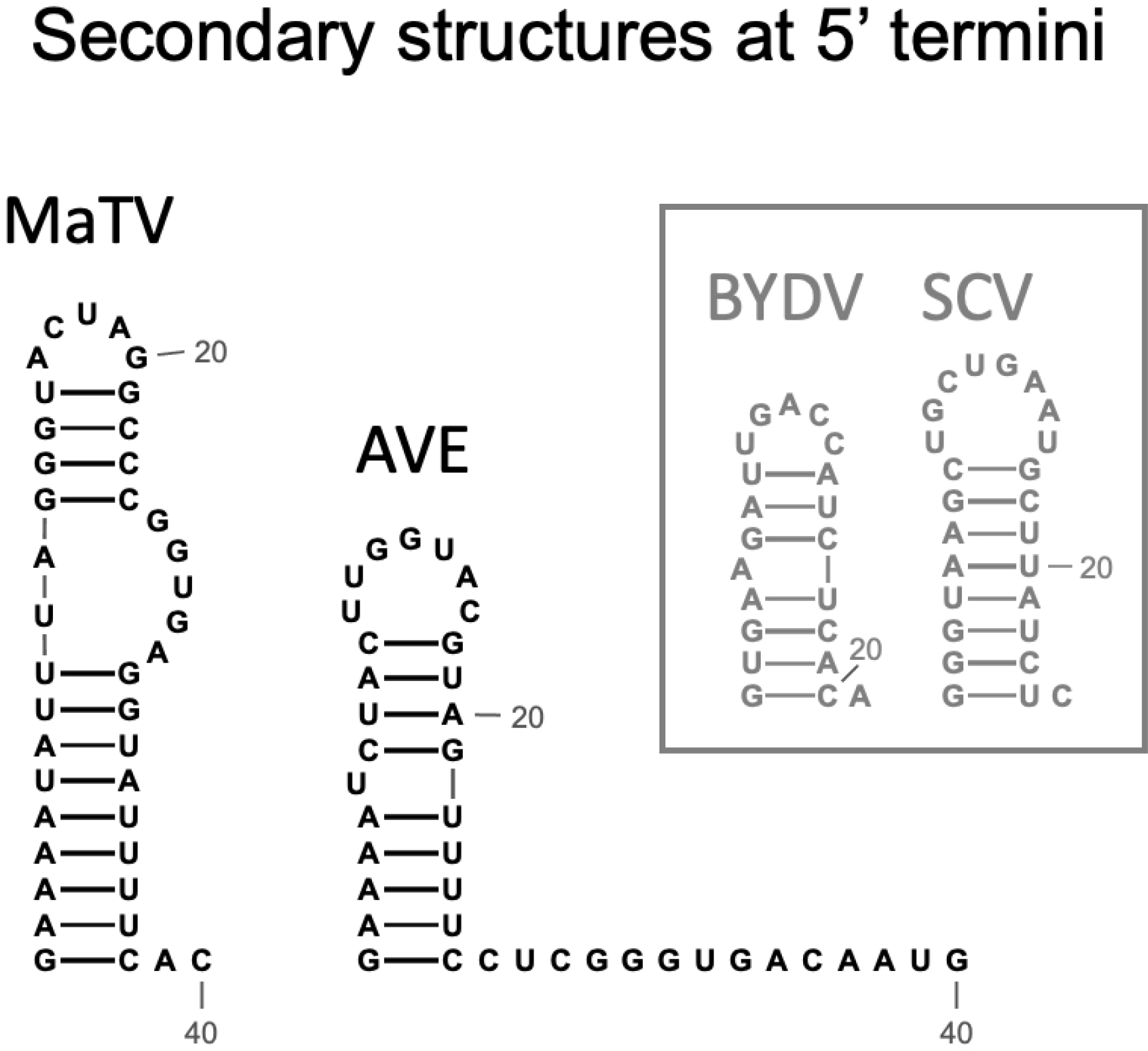
Predicted secondary structures of the 5’ termini of MaTV and AVE, the only two rimosaviruses for which the 5’ end is known (MaTV) or predicted (AVE). For comparison, secondary structures determined by chemical probing of tombusvirids barley yellow dwarf virus (BYDV, *Luteovirus* (Guo et al., 2001)), and saguaro cactus virus (SGV, *Carmovirus* (Chattopadhyay et al., 2011)) are shown.

We predict that the sequence upstream of ORF1 is untranslated in rimosaviruses based on the following observations. (i) There are no conserved ORFs upstream of ORF1. MaTV contains a predicted AUG-initiating ORF of 336 nt (112 codons), that we call ORF0, starting at nt 85, but it is absent in the other rimosaviruses, save for a truncated version of ORF0 (177 nt, 58 codons) in PVLaTV1 (Fig. 1), the closest relative to MaTV (75.4% nt identity). (ii) There are 6 to 12 AUG triplets scattered in different positions among the rimosavirus genomes upstream of the ORF1 start codon (Table 2, Supplementary Fig. S1), including upstream and downstream of ORF0 in MaTV and PVLaTV1, but they result in short, non-conserved ORFs. (iii) Finally, ORF1 is the 5’-proximal ORF in all other tombusvirids.

**Table 2.** Properties of rimosavirus genomes.

| virus | length of sequenced genome<sup>a</sup> | uAUGs<sup>b</sup> | Gs/Cs in 24 nt 5' of ORF1 | 5' UTR length<sup>c</sup> | 3' UTR length | Approx. 3' nt absent<sup>d</sup> | In-frame ORF4 AUG<sup>g</sup> | (PX)n repeats in RTD<sup>h</sup> |
| --- | --- | --- | --- | --- | --- | --- | --- | --- |
| MaTV | 5315 nt | 12 | 0G/10C | 622 nt | 434 nt | 0 | yes | 4 + 6 |
| PVLaTV1 | 5110 nt | 9 | 1G/7C | 464 nt | 418 nt | 15 | no | 2 + 5 |
| AVE | 5398 nt | 11 | 0G/12C | 597 nt | 458nt | 13 | no | 0 |
| TTTV1 | 5245 nt | 8 | 0G/9C | 662 nt | 374 nt | 32 | yes | 3 |
| ENaTV10 | 5293 nt | 8 | 1G/6C | 658 nt | 425 nt | -21<sup>e</sup> | no | 3 |
| NTTV1 | 5461 nt | 6 | 0G/11C | 641 nt | 412 nt | ? | yes | 2 |
| HubTLV2 | 5510 nt | 7 | 1G/14C | 562 nt | 437 nt | 2? | yes | 3 |
| BCaTV | 5447 nt | 8 | 0G/9C | 727 nt | 229 nt | ?<sup>i</sup> | no | 7 |
| ZLaTV | 5255 nt | 6 | 0G/9C | 457 nt | 247 nt | ?<sup>i</sup> | no | 0 |
| HubTLV1 | 5430 nt | 9 | 0G/11C | 515 nt | 435 nt | ~6-8 | no | 2 + 3 + 2 |
| ENaTV5 | 5631 nt | 12 | 0G/14C | 438 nt | 474 nt | ~6-8 | no | 0 |
| TCaTV1 | 5501 nt | 10 | 0G/13C | 680 nt | 410 nt | ~27-29 | no | 2 |

<sup>a</sup>Sequence of Genbank accession numbers in Table 1. <sup>b</sup>AUGs upstream of ORF1 start codon. <sup>c</sup>5' UTR is defined as sequence upstream of ORF1 start codon. All but MaTV and AVE are likely incomplete. <sup>d</sup>Nts missing estimated by alignments of sequences and secondary structures from the conserved GGCG-bulged stem-loop to the 3' end of Genbank sequence (Fig. 5, Supplementary Fig. S3). <sup>e</sup>Appears to have nt beyond the 3' terminal GCCC. <sup>f</sup>3' UTR sequence is too incomplete to have GGCG bulge. <sup>g</sup>AUG start codon. <sup>h</sup>n = number of consecutive alternating proline repeats [(PX)n] in the readthrough domain (RTD) encoded by the first 200 nt of ORF5 (3'-adjacent to ORF3 stop codon) (P = proline, X = any amino acid). Plus signs indicate separate blocks of (PX)n repeats.

Such a 5’ UTR consisting of many hundreds of bases (Table 2) and containing numerous nontranslated AUG triplets suggests presence of an internal ribosome entry site (IRES) or possibly a long ribosomal shunt system. IRESes in the genomes of picornaviruses, hepaciviruses and many other animal RNA viruses are highly structured RNAs that co-opt various host proteins to recruit the ribosome shortly upstream of the first actual start codon for highly efficient cap-independent translation (Filbin and Kieft, 2009;Fraser et al., 2009;Yamamoto et al., 2017). In shunting, scanning ribosomes jump across a long structured stem-loop, then rejoin the mRNA to scan to the start codon (Ryabova et al., 2002). However, we detect little sequence conservation among the 5’ UTRs of the twelve rimosaviruses, nor were we able to predict any conserved secondary structure, as would be expected for an IRES or a shunting mechanism. One conserved feature is that the sequence upstream of and near the ORF1 AUG is G-poor and C-rich (Fig. 4). Specifically, in all twelve viruses, the 24 nt immediately upstream of ORF1 start codon have at most one guanosine base (Table 2). The C-richness usually extends 50 or more bases upstream of the ORF1 AUG (Fig. 4). Certain IRESes have cytosine- or pyrimidine-tracts at about this position relative to the start codon (Jaramillo-Mesa et al., 2018), and the region just upstream of well-translated start codons in plants is enriched in C-rich tracts (Wu et al., 2024). In summary, the 5’ UTR is enigmatic, as it contains numerous AUG codons and small nonconserved ORFs, that seem to rule out a conventional scanning mechanism for initiation of translation of ORF1, but it appears to lack conserved secondary structures or sequences (except potential C-rich motif), as are known in IRESes or shunting structures.

**Fig. 4.**
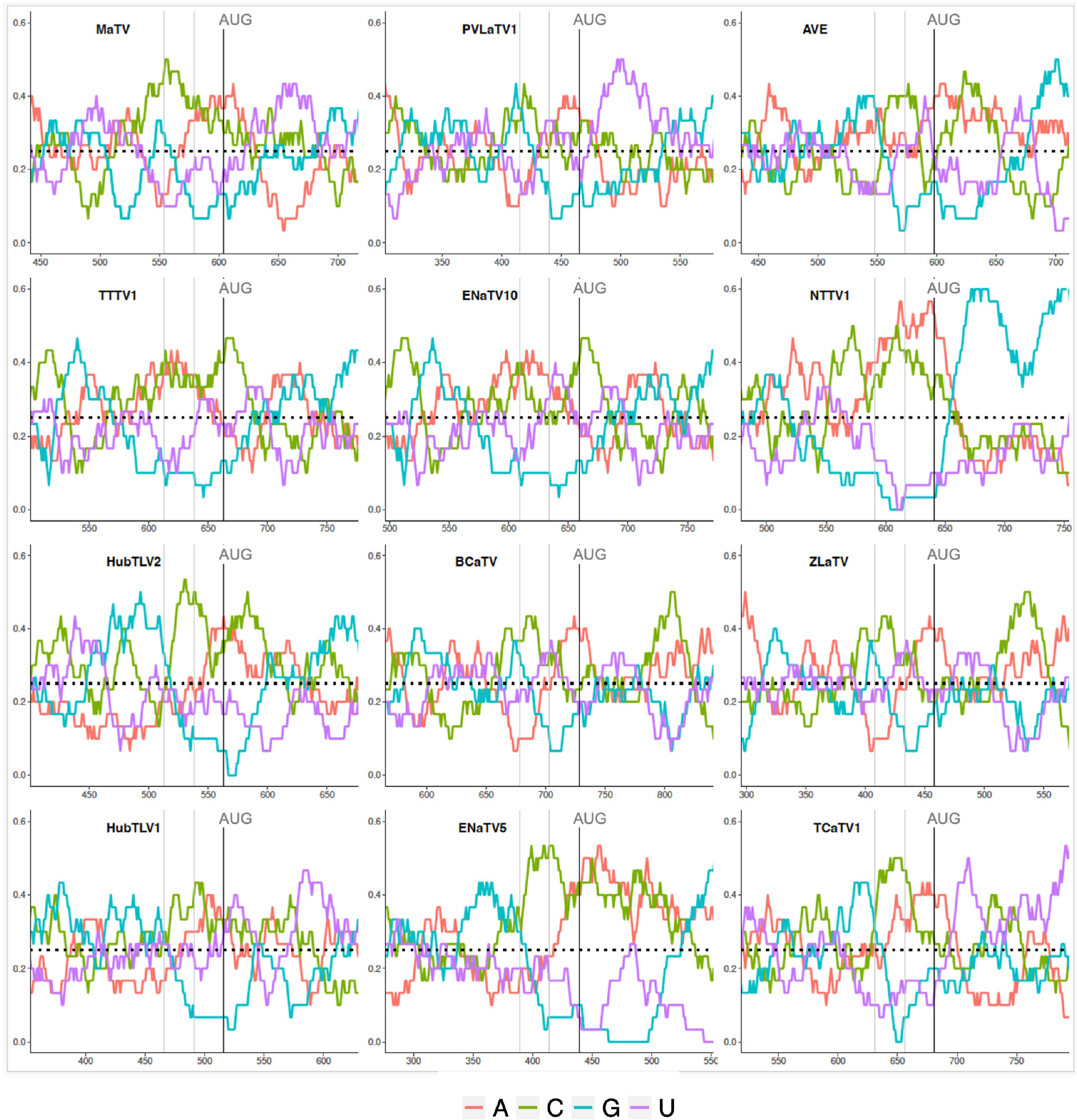
Base compositions flanking the ORF1 start codons (AUG). Plots depicting relative nucleotide frequencies calculated using a sliding window approach plus or minus one-hundred fifty or one-hundred nucleotides relative to the ORF1 start codon, respectively. The X-axis shows genomic position, the Y-axis represents the frequency of each base in a 50 nucleotide window as depicted by the colors in the legend.

#### 3.2.2 3’ untranslated region

The 3’ terminus of MaTV is CCC, the trinucleotide present at the 3’ end of virtually all tombusvirid genomes for which the 3’ sequence has been confirmed. The eleven posted rimosavirus sequences other than that of MaTV do not end in CCC, so we predict they are incomplete. To estimate roughly how many bases are missing from the 3’ ends, we took advantage of known structures near the 3’ end. Tombusvirids have a distinct conserved secondary structure at the 3’ end in which the final four bases, usually GCCC, form a pseudoknot by base pairing to a GGGC bulge in a nearby upstream stem-loop (Koev et al., 2002;Pogany et al., 2003;Na et al., 2006;McCormack et al., 2008;Simon, 2015). Indeed, we predict these structures in MaTV RNA, with the bulged stem-loop terminating 61 nt upstream of the 3’ end of the genome (Fig. 5A).

**Fig. 5.**
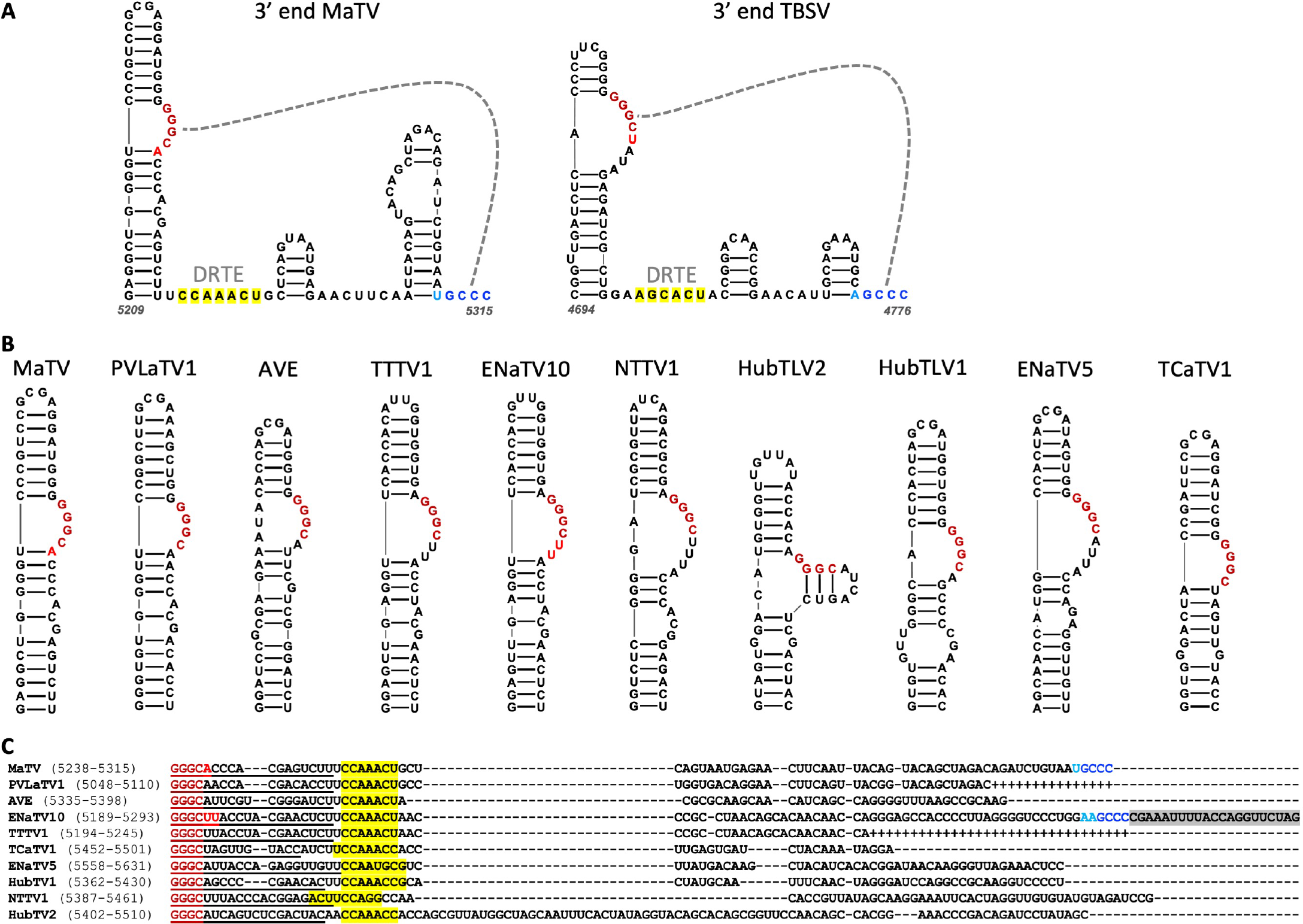
A. Predicted secondary structure using Mfold and Scanfold of the 3’-terminal 107 nt of the MaTV genome, and the known secondary structure of the 3’ end of the TBSV (*Tombusvirus*) genome (Na and White, 2006;Cimino et al., 2011). Gray dashed line indicates pseudoknot base pairing. Dark red and blue bases indicate conserved GGGC:GCCC base pairing, while lighter red and blue indicate additional base complementarity. Yellow highlighting indicates bases in the putative distal readthrough element (DRTE) predicted to pair to the bulged stem-loop adjacent to ORF1 stop codon (proximal readthrough element, PRTE, Fig. 9). **B.** GGGC-bulge-containing stem-loops upstream of 3’ terminus of genome as found in other tombusvirids.. BCaTV and ZLaTV are not shown because their GenBank sequences presumably terminate upstream of the GGGC bulge. **C.** Alignments of 3’ ends (MUSCLE) of available 3’ terminal sequences starting at the GGGC bulge (dark red). Known (MaTV) and predicted (ENaTV10) 3’ terminal GCCC are in blue (see also Supplementary Fig. S3). Additional potential base pairing between GGGC bulge region and 3’ end are shown in lighter shades of red and blue, respectively. Plus symbols in place of dashes indicate missing bases based on closely related sequence above the sequence. Predicted distal readthrough element (DRTE) capable of base pairing to a bulge in the proximal readthrough element (PRTE, Fig. 9) is highlighted in yellow. Predicted nonviral sequencing adapter-derived sequence is highlighted in gray (ENaTV10). Numbers in parentheses indicate base positions in the available genome sequence.

To estimate how many bases are missing from the 3’ ends of the other rimosaviruses, we searched for the 3’-proximal GGGC tract in a bulged stem-loop in each genome. All but two viral genomes gave a discrete stem-loop with the GGGC bulge (Fig. 5B). (Although weak intra-bulge base pairs were predicted in HubTLV2). The two exceptions, BCaTV and ZLaTV, have much shorter 3’ UTRs in the GenBank sequences (229 nt and 247 nt, respectively) than the others, which range from 374 to 474 nt (Table 2); thus the posted BCaTV and ZLaTV 3’ UTR sequences are both probably incomplete to the extent that they lack the entire GGGC bulged stem-loop. Based on the number of bases downstream of this bulged stem-loop in the other rimosavirus sequences, we roughly estimated the number of bases that would be missing from each 3’ end if the number of bases from the bulged stem-loop to the 3’ end is the same as for MaTV (Table 2). However, the HubTLV2 and ENaTV10 3’UTRs extend beyond the predicted position for the terminal GCCC sequence and do not terminate in GCCC (Fig. 5C). An additional stem-loop adjacent to the terminal GCCC is much larger than that of MaTV, giving HubTLV2 and ENaTV10 longer 3’UTRs (Supplementary Fig. S3). ENaTV10 has an additional 21 nt 3’ of a GCCC tract. We speculate that adapter bases used for sequencing were not trimmed from what is the true 3’ end of the viral genome (GCCC). Interestingly, the predicted terminal six bases, AAGCCC can form a six-base pseudoknot with the upstream GGGCUU bulge, instead of the usual four base pairs (Supplementary Fig. S3).

Tombusvirid RNAs are uncapped (Allen et al., 1999), so they carry a cap-independent translation element (CITE) in the 5’ end of the 3’ untranslated region (Simon and Miller, 2013). In this region of the rimosavirus genomes we did not identify any secondary structures conserved across all twelve viruses. We predict structures conserved between closely related viruses MaTV and PVLaTV1, and between ENaTV10 and TTTV1 (Supplementary Fig. S4). However, these two pairs of structures differ from each other and do not resemble known CITEs. That said, CITEs can be difficult to identify, given the variety of structures possible, varying from a bulged stem-loop with some conserved motifs to more complex branched structures (Mizumoto et al., 2003;Fabian and White, 2006;Wang et al., 2011;Nicholson et al., 2013). Thus, we cannot rule out the presence of a 3’ CITE in the 3’ UTRs of rimosaviruses, given that the 3’ UTRs are certainly long enough to contain a 3’ CITE.

### 3.3 Rimosavirus ORFs and encoded proteins

#### 3.3.1 ORFs 1-2

ORF1 is predicted to be translated as the protein P1 and also fused with ORF2, via occasional stop codon readthrough, to produce P1-P2 protein. As expected, P2 contains the RdRp active site with the highly conserved D(X)3<D, SG(X)3T(X)3N(X)25GDD motifs (Charon et al., 2022). In addition to forming a clade distinct from those of other tombusvirids (Fig. 2), the RdRps also fall into two subclades with MaTV, PVLaTV1, ENaTV10, TTTV and AVE in one group. The other seven rimosaviruses contain a 67 to 81 amino acid insertion between the positions of amino acids 85 and 86 in the RdRps of MaTV, PVLaTV1, ENaTV10, TTTV and AVE (Supplementary Fig. S2).

#### 3.3.2 ORF3

ORF3 encodes the coat protein, as evidenced by homology with those of other tombusvirids. The CPs of the twelve rimosaviruses fall into a distinct clade like the RdRps, but some relationships within the proposed genus differ from those of the RdRps (Fig. 6). For example, the CPs of TTTV1/ENaTV10 are in a distant branch from MaTV and PLVaTV1, but the RdRps of these four viruses fall in the same subclade (Fig. 2). While the RdRps of BCaTV and ZLaTV are closely related, their CPs are not. Interestingly the CPs of other tombusvirids – with the exception of luteoviruses - are more closely related to those of viruses in genus *Sobemovirus* of the *Solemoviridae* family (e.g. rice yellow mottle virus, RYMV), than they are to those of the rimosaviruses (Fig. 6).

**Fig. 6.**
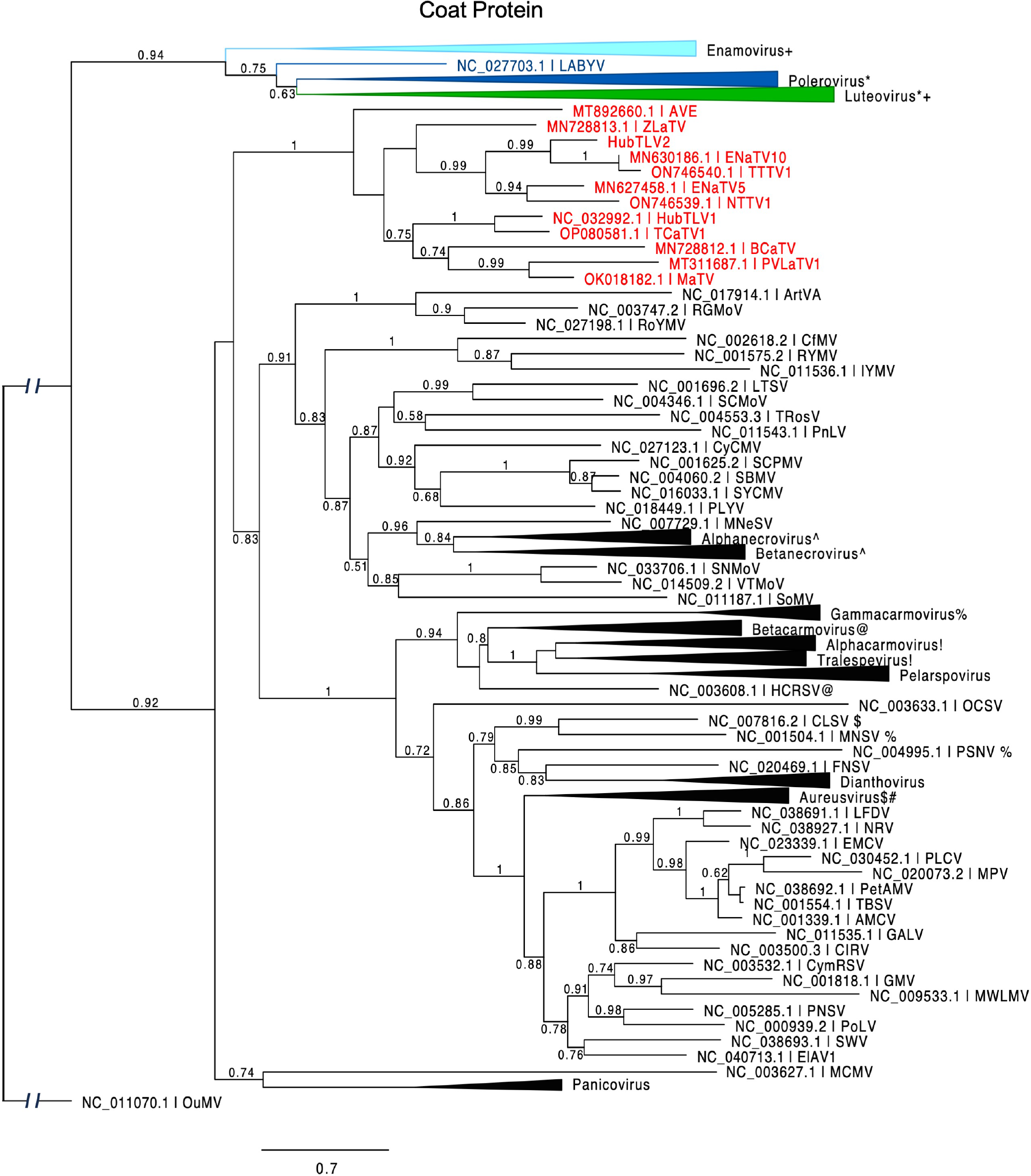
Phylogenetic tree predicting the relationship of viral coat proteins (ORF3) in *Tombusviridae* and *Solemoviridae*. Red entries indicate those members of the proposed new *Rimosavirus* genus. Each collapsed tree consists primarily of sequences belonging to the indicated genus. Modifier symbols (+, *, ^, %, etc.) indicate genera where one or more CP sequences from one genus are grouped with the other genus containing same modifier symbol. Branch support values are shown for splits > 0.5 and are calculated from 1,000 resamples of the Shimodaira-Hasegawa test (SH-like local supports). Branch lengths indicate arbitrary units of evolutionary distances. Spell-outs of viral acronym can be found in GenBank via the indicated accession number. Because solemovirids (*Pisuviricota*) and tombusvirids (*Kitrinoviricota*) belong to separate phyla, the outgroup CP sequence (Ourmia melon virus, OuMV) was chosen from a third phylum, *Lenarviricota*.

#### 3.3.2 ORF4

Poleroviruses and most luteoviruses (but not enamoviruses) encode a movement protein (MP) in ORF4, that overlaps with ORF3, initiating a few nucleotides downstream of the ORF3 start codon but in a different frame and terminating shortly before the ORF3 stop codon (Domier, 2012). In these genera, ORF4 is translated from the same subgenomic mRNA (sgRNA1) as ORF3 (Koev et al., 1999;Juszczuk et al., 2000), via leaky scanning, in which some scanning ribosomes skip the ORF3 AUG and instead initiate on the ORF4 AUG codon (Dinesh-Kumar and Miller, 1993;Miras et al., 2017). In contrast, in four of the rimosaviruses, what we call ORF4 appears to initiate with an AUG codon upstream of the ORF3 start codon. In the other eight, there is a similar ORF overlapping with most of ORF3, but it lacks an in-frame methionine start codon and appears to be disrupted by frameshift mutations upstream of the ORF3 overlap (Fig. 7). Upon aligning the RNA sequence at the beginning of MaTV ORF4 with the homologous PVLaTV1 sequence, we see an AUG in PVLaTV1 sequence that aligns with the second AUG of MaTV ORF4 as does the remaining sequence until position 48 at which a U insertion occurs in PVLaTV1 RNA (Fig. 7B). This places the upstream portion of what would encode PVLaTV1 ORF4 out of frame with the rest of the ORF. A U insertion at a similar position also disrupts ORF4 in ENaTV10 ORF4 relative to the 95% identical TTTV1 genome, which has an intact AUG-initiated ORF4 (Fig. 7B). Similar frameshift mutations near the 5’ end of ORF4 may explain why the ORFs 4 of AVE, BCaTV, ZLaTV, HubTLV1, ENaTV5 and TCaTV1 also appear to lack an in-frame AUG start codon (Supplementary Fig. S5).

**Fig. 7.**
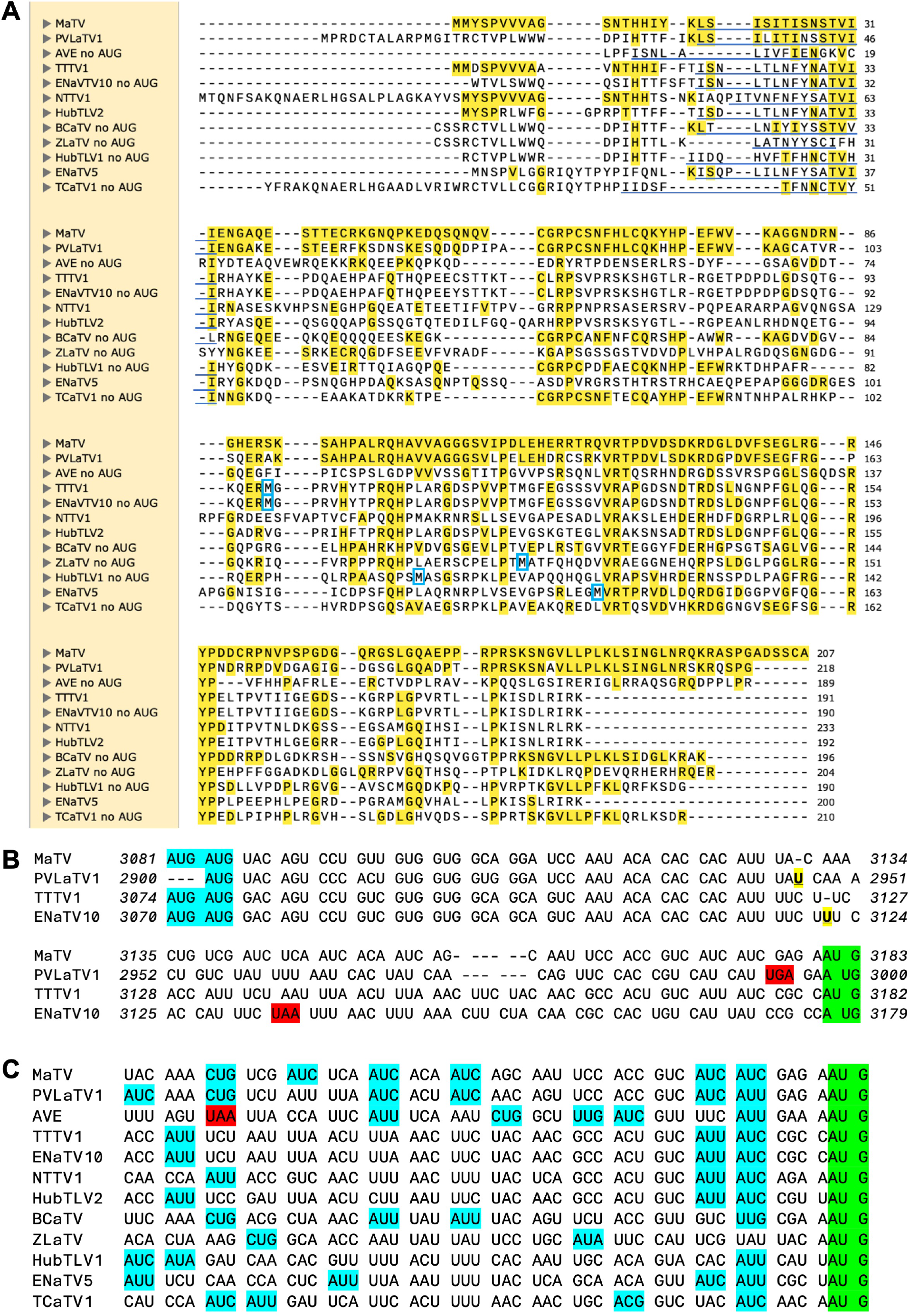
ORF4 comparisons. **A.** Alignment (MUSCLE) of predicted P4 proteins, starting from an in-frame methionine or immediately downstream of an in-frame stop codon. However, actual predicted N-terminus of P4 is predicted to be from translation initiation at a non-AUG codon in the underlined region (per ungapped nt alignment, panel C). First met translated from an AUG located downstream (in the ORF4 frame) of the ORF3 AUG is indicate by blue box. MaTV, PVLaTV1, AVE, HubTLV2, BCaTV and TCaTV1 have no such AUGs (i.e. no internal mets). Yellow highlighting: aa identical to that of MaTV P4. **B.** Alignment of the regions of ORF4 from their potential AUG start codons (blue) to the ORF3 start codons in a different frame (green), with spacing to show codons starting from the AUGs. The U insertions (bold, underlined, yellow) in PVLaTV1 relative to MaTV, and in ENaTV10 relative to TTTV1, disrupt the reading frames, leading to stop codons (red). For PVaLTV1 and ENaTV10, the potential initiator AUGs here in a different frame than the amino termi in panel A. Italic numbering indicates genomic positions of first and last nucleotide on each line. See Supplementary Fig. S5 for alignments of this portion of all 12 rimosavirus genomes without codon spacing. **C.** Possible non-AUG start codons (blue) (Gordon et al., 1992;Diaz de Arce et al., 2018;Fang and Liu, 2023) for ORF4 are shown in the 52 nt upstream of the ORF3 start codon (green). Stop codon is highlighted in red.

Other possibilities were considered. If the 5’ end of the sgRNA1 is downstream of the predicted ORF4 AUG start codon but upstream of the ORF3 start codon, then we would expect that ORF4 would initiate at the next AUG downstream of the ORF3 start codon, as in the luteo- and poleroviruses (Dinesh-Kumar and Miller, 1993). However, AVE, BCaTV1, TCaTV1 and PVLaTV1 have no AUG codons anywhere in ORF4, and ORFs 4 of MaTV and HubTLV2 have no AUG codons downstream of the ORF3 start codon (methionine residues, Fig. 7A). The AUG codons in ORFs 4 of the other rimosaviruses are in the middle or C-terminus of the ORF and not in conserved locations (Fig. 7A). Thus, initiation by leaky scanning downstream of the ORF3 start codon is highly unlikely. Instead, the most plausible explanation is that ORF4 initiates with a non-AUG codon. The non-AUG codons ACG, AUC, AUU, AUA, UUG, CUG, GUG have been observed to serve as start codons, albeit much less efficiently than AUG (Gordon et al., 1992;Diaz de Arce et al., 2018;Fang and Liu, 2023). To allow translation of ORF4 in all twelve rimosaviruses, a non-AUG start codon would have to be located downstream of the homologous positions to the frame-disrupting insertions in ENaTV10 and PVLaTV1 and an in-frame stop codon in AVE (Fig. 7C), and upstream of the ORF3 AUG start codon, as initiation of translation at a non-AUG is not likely to take place downstream of the highly efficient AUG. Alignment of this portion of the rimosavirus genome revealed numerous non-AUG codons, known to be capable of initiation (Fig. 7C). Thus, we propose that ORF4 is translated by initiation at one of these non-AUG start codons. This arrangement allows leaky scanning initiation at the AUG of ORF3, resembling the arrangement of ORF3a in luteo- and poleroviruses, which also initiates with a non-AUG codon and overlaps with ORF3 (Smirnova et al., 2015).

#### 3.3.3 ORF5

The arrangement of rimosavirus ORFs 3 and 5 resembles that in the L/P/E genera because ORF5 appears to be translated by readthrough of the ORF3 stop codon, creating a large C-terminal readthrough domain (RTD) extension to the CP. The L/P/E RTDs all are more closely related to each other than to any of those from proposed genus *Rimosavirus*, despite the fact that luteoviruses (*Tombusviridae*) and polero- and enamoviruses (both *Solemoviridae*) belong to different families. Interestingly, polerovirus RTDs (dark blue in Fig. 8) fall into two major subclades. One clade also includes a luteovirus (green) RTD (bean leafroll virus, BLRV), while the other polerovirus clade includes enamovirus RTDs (light blue). The RTDs of the rimosaviruses (red) all diverge highly from those of L/P/E viruses and from each other. Although some bootstrap values are low, RTDs of the branches representing HubTLV2/ENaTV10/TTTV1, MaTV/PLVaTV1, AVE, and ZLaTV all are more divergent from each other than all of the divergence among all the L/P/E viruses.

**Fig. 8.**
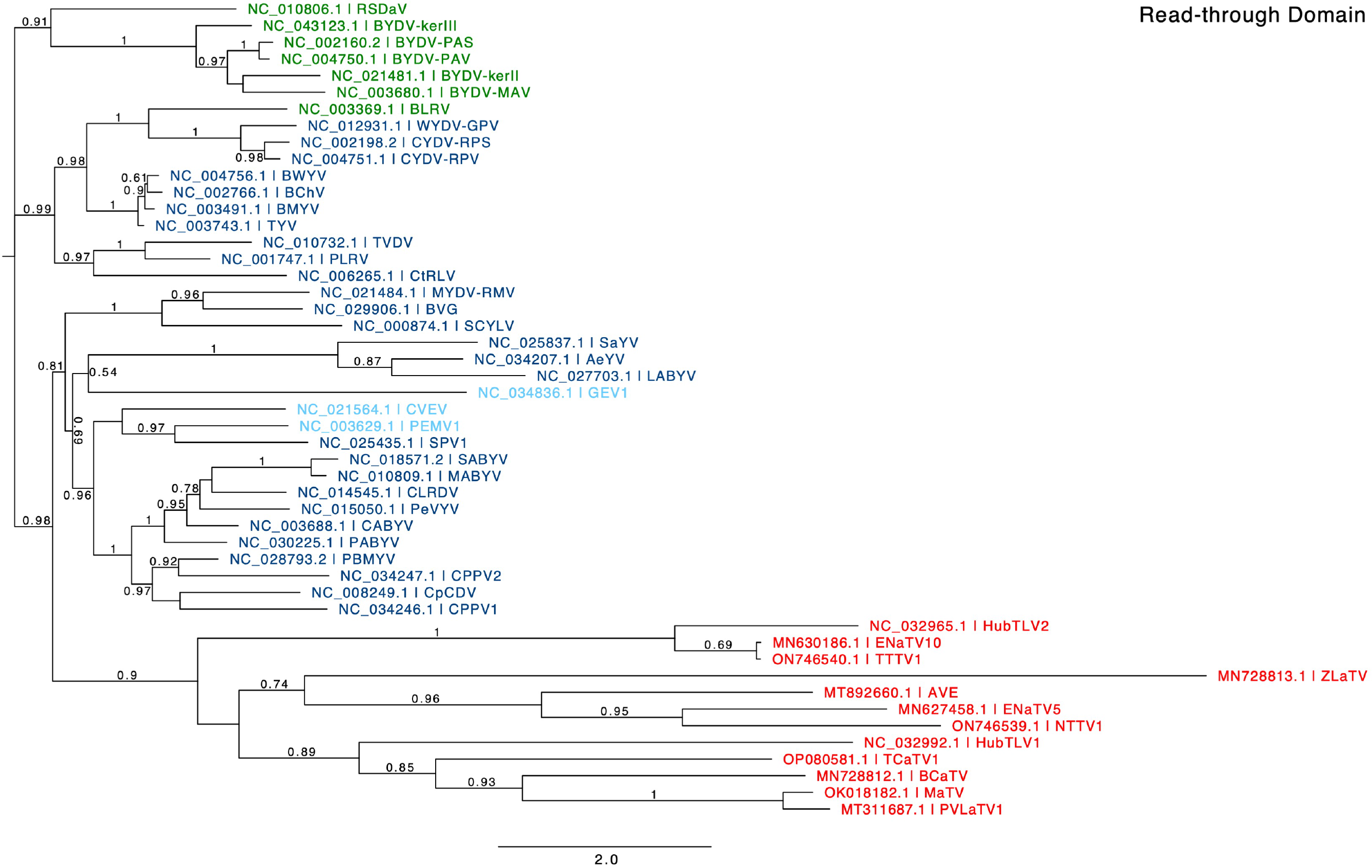
Phylogenetic tree predicting the relationships of RTDs (ORF5) based on the amino acid sequences. Genus is color coded in red for *Rimosavirus*, green for *Luteovirus*, dark blue for *Polerovirus* and light blue for *Enamovirus*. Branch support values are shown for splits > 0.5 and are calculated from 1,000 resamples of the Shimodaira-Hasegawa test (SH-like local supports). Branch lengths indicate arbitrary units of evolutionary distances. Full virus names can be looked up via the indicated GenBank accession number.

Features of ORF5 and the RTD it encodes reflect these extreme differences from the L/P/E RTDs. For example, ORF5 of L/P/E virus RNAs contains eight to sixteen direct repeats of the sequence CCXXXX (X = any base) shortly downstream of the ORF3 stop codon. This encodes an alternating proline repeat (PX)n, which is thought to serve as a spacer between the CP and functional domains of the RTD protein (Mutterer et al., 1999;Bonning et al., 2014;Schiltz et al., 2022). However, most rimosaviruses have few, if any CCXXXX or PX repeats in the RNA and encoded protein, respectively (Table 2).

### 3.4 Readthrough elements

Readthrough of stop codons in viral RNAs is usually facilitated by RNA structures located immediately 3’ of the stop codon (Brown et al., 1996;Cimino et al., 2011;Newburn et al., 2014;Newburn and White, 2017;Xu et al., 2018b;Chkuaseli and White, 2022). The ∼100 nt adjacent to the ORF1 stop codons of all twelve rimosaviruses are well-conserved in sequence and secondary structure (Fig. 9). This UAG-proximal structure consists of a stem-loop with four to five helices separated by bulged regions, including a distal bulge with a run of 3 C’s, and a more proximal bulge with the consensus RGUUUGG (red, Fig. 9). We predict this conserved sequence base pairs with downstream sequences to form a pseudoknotted structure that facilitates readthrough, as shown for other tombusvirids (Cimino et al., 2011;Chkuaseli and White, 2018). Indeed, in the 3’ UTR, just downstream of the GGGC bulged stem-loop is a conserved CCAAAYY sequence in a region predicted to be single stranded (Fig. 5C, Supplementary Fig. S3). This is the exact position of the distal readthrough element (DRTE) which base pairs to a bulge in the stop codon-proximal readthrough element (PRTE) to facilitate readthrough in CIRV (genus *Tombusvirus*) (Cimino et al., 2011). A DRTE is also at or near this position in the genomes of at least six other tombusvirid genera, all base pairing to the PRTE with different sequences (Cimino et al., 2011).

**Fig. 9.**
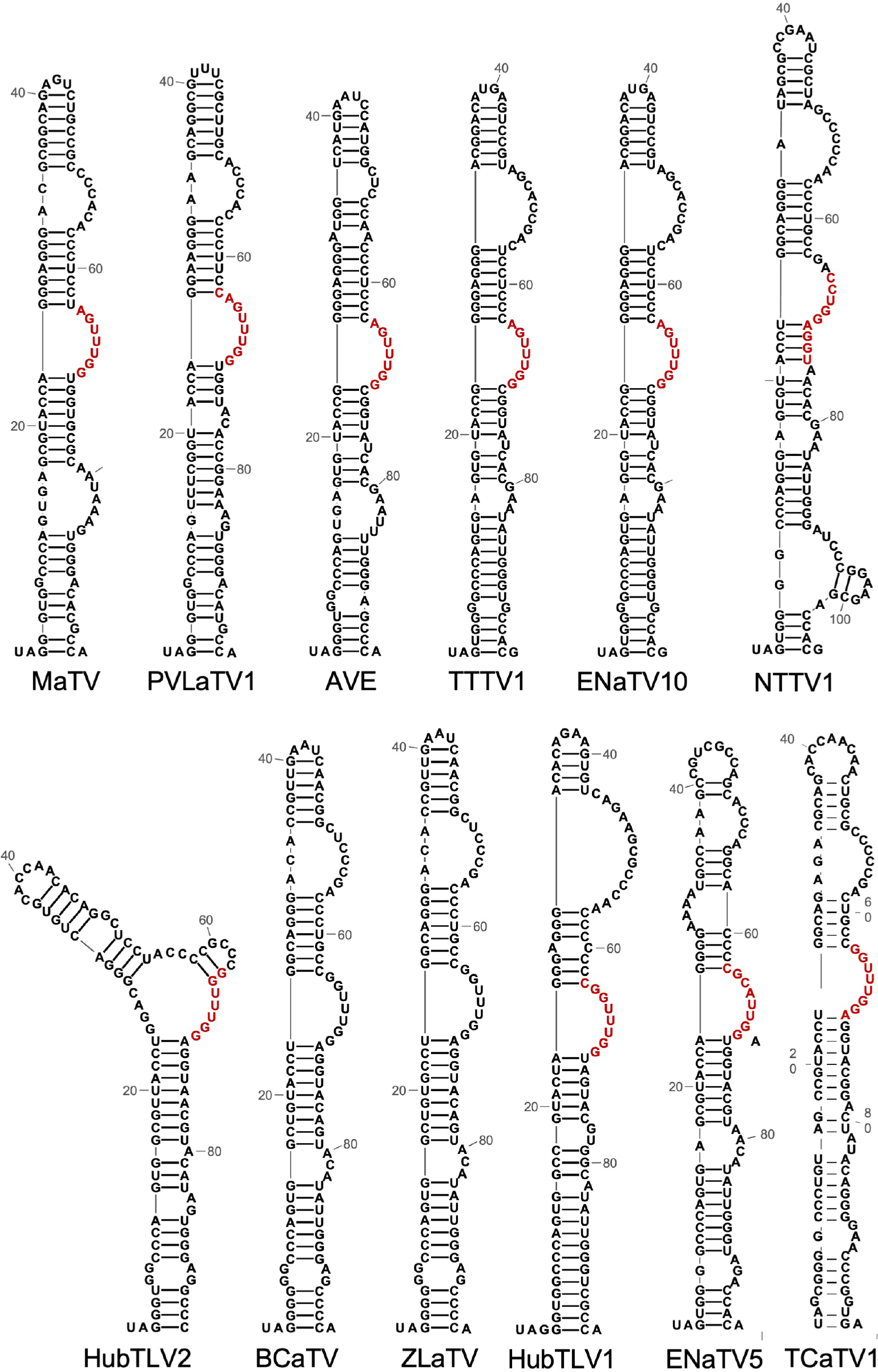
Predicted secondary structures (RNA Alifold, Mfold) of a sequences beginning with the putatively leaky ORF1 stop codon (UAG in all rimosaviruses). Based on studies of other tombusvirids, these structures comprise the proximal readthrough element (PRTE). Bases in red are predicted to base pair to the distal readthrough element (DRTE, highlighted in yellow in Fig. 5). Note the co-variations in which the sequences vary but maintain at least seven consecutive base pairs in all 10 viruses where the base pairing can be predicted. This long-distance base pairing cannot be predicted for BCaTV and ZlaTV PRTEs because the available genome sequences lack the region containing the PRTE.

For readthrough of the predicted leaky ORF3 stop codon, we look to research on RNA sequences and structures that control readthrough of the homologous stop codon in L/P/E viruses. Mutagenesis of the (CCXXXX)8-16 repeat sequence prevented efficient readthrough for BYDV (Brown et al., 1996) and PLRV (Xu et al., 2018b). Andy White’s lab then did a more comprehensive analysis of the Pea enation mosaic virus 1 (PEMV1) (*Enamovirus*) readthrough structure which revealed four separate long-distance base pairings that coaxially stack in the readthrough-facilitating structure (Chkuaseli and White, 2022). That publication also revealed a different arrangement of long-distance base pairings needed for PLRV readthrough. Finally, they showed that local base pairing can compete with the long-distance base pairing to perhaps comprise a switch to regulate readthrough efficiency (Chkuaseli and White, 2022). Because the RTDs of the rimosaviruses are so divergent (Fig. 8), there was not enough sequence similarity to identify a consensus secondary structure. We predict diverse stem-loop structures adjacent to the ORF5 stop codon, but we found no common secondary structures (Supplementary Fig. S6). Thus, the RTD for rimosaviruses is novel, not only in its high amino acid sequence variation in its encoded RTD, but also for lack of obvious conserved sequence (e.g. CCXXXX repeats) or obvious L/P/E-like secondary structure to facilitate the CP ORF stop codon readthrough.

## 4 **Discussion**

### 4.1 Remarkable distribution and diversity of sources of rimosaviruses

Rimosavirus sequences were collected from diverse organisms around the globe in large environmental metagenomics sequencing projects with no reports of actual hosts in which they replicate or disease symptoms they cause. Because of their worldwide distribution, it is perhaps surprising that the viruses associated with these genomes have not been discovered previously. It is remarkable that the genome of the virus we call MaTV because its genome was found first in maize and its ancestor teosinte, in Mexico, was also found in the cloaca of a tuatara on the tiny uninhabited island of Takpourewa (also known as Stephens Island) in New Zealand (Table 1), a wildlife refuge on which no crops are cultivated (East et al., 1995). Moreover, BCaTV and ZLaTV sequences, first described in kohlrabi and Manchurian wild rice, respectively, in China (Yang et al., 2022) were also found in the Asian long-horned tick in China (Ni et al, 2023) and in the cloaca of a tuatara in New Zealand (Waller et al., 2022). These wide distributions suggest these viruses may have wide host ranges, and perhaps rather cryptic symptomatology. However, the host range is likely limited to plants, based on their clear membership in the *Tombusviridae* family, despite the fact that several were isolated from various invertebrates, a reptile, and from plant pathogenic fungi (Table 1). The rimosaviruses associated with *Erisyphe necator* (powdery mildew of grapevine) and *Plasmopara viticola* (downy mildew of grapevine) may have infected plant material contaminating the mildew preparation for sequencing, or the mildew may be a vector of the virus. The tombusvirid cucumber necrosis virus is transmitted by zoospores of the soil fungus *Olpidium bornovanus* (Rochon et al., 2004). Recently, cucumber mosaic virus (not a tombusvirid), which has a wide plant host range, was shown to infect and replicate in the plant pathogenic fungus *Rhizoctonia solani* (Andika et al., 2017). Thus, we cannot rule out that these apparent plant viruses may infect the mildews with which they associate.

The animal-associated rimosavirus genomes may have been acquired from the plant material in the diet of Chinese land snails (HubTLV1) or pill worms (HubTLV2). For the carnivorous tuatara, the rimosavirus found in its cloaca could be derived from a herbivorous insect in its diet. Plant viruses, including tombusvirids, have been found in other carnivores such as dragonflies (Li et al., 2010) and bats (Li et al., 2010), and were assumed to have been obtained this way. Ticks are blood feeders, but even in that case, plant viral sequences, including those of tombusvirids have been identified in the human blood virome, albeit at very low abundance (Cebriá-Mendoza et al., 2021). This wide diversity of associations by rimosaviruses is a testament to the high abundance and particle stability of tombusvirids. Clearly, additional experiments are necessary to determine the actual hosts in which these rimosaviruses replicate.

### 4.2 Gene organization and protein function

As mentioned above, the rimosavirus genome encodes P1 and P2 almost certainly via a readthrough mechanism, which is standard for most tombusvirids. In TBSV, and probably all tombusvirids, P1 binds viral RNA and lines the membrane-bound replication vesicles, while P1-P2 fusion has the RNA-dependent RNA polymerase activity to replicate the genome and transcribe subgenomic mRNA (Nagy, 2016). ORF3 clearly encodes the CP to form the T=3 icosahedral virion, based on sequence similarity to other tombusvirids.

We expect P4 plays a role in virus movement and other functions, based on the fact that ORF4, which also overlaps with ORF3 in polero- and luteoviruses, encodes a movement protein (MP) in these viruses (Chay et al., 1996;Schmitz et al., 1997;Link et al., 2011). It can also support virus infection by suppressing (i) antiviral RNA silencing (Fusaro et al., 2017), (ii) host catalase activity (Tian et al., 2024), and (iii) thiamine synthesis (Han et al., 2024). ORF4 differs in that we predict its translation initiates upstream instead of downstream of the ORF3 start codon, and at a non-AUG start codon. Thirdly, we detected no ORF3a, which encodes another MP in the ORF4- encoding viruses (Smirnova et al., 2015;DeBlasio et al., 2018). Thus, translationally, rimosavirus ORF4 is like a fusion of luteo-/polerovirus ORFs 3a and 4.

While unlikely, it cannot be ruled out that ORF4 may not be translated in the rimosaviruses that lack an in-frame AUG start codon for ORF4. Recently some luteoviruses were discovered that lack ORFs 4 and 3a (Khalili et al., 2023;Stainton et al., 2023). None of the polero-like enamoviruses encode ORFs 3a or 4 (Smirnova et al., 2015). For some enamoviruses, the movement functions are provided by a co-infecting umbravirus (Ryabov et al., 2001a;Ryabov et al., 2001b), but for other enamoviruses and the luteoviruses lacking ORFs 3a and 4, no co-infecting partner is known. Thus these and many rimosaviruses may have found other ways to move within the host plant, perhaps by commandeering a host phloem protein as has been observed recently for certain umbravirus-like viruses in the *Tombusviridae* (Ying et al., 2024). However, given the presence of ORF4 overlapping with most of ORF3 in all twelve rimosavirus genomes, we favor our hypothesis that ORF4 is translated via initiation at a non-AUG shortly upstream of the ORF3 AUG start codon.

Based on its position and homology to ORF5 in L/P/E viruses, ORF5 is highly likely to be translated by readthrough of the ORF3 stop codon, and thus encodes the RTD. In the L/P/E/s, the RTD is essential for the persistent, circulative, nonreplicative transmission by aphids (Chay et al., 1996;Gunasinghe et al., 1997;Brault et al., 2000;Gray and Gildow, 2003;Schiltz et al., 2022). Thus, we speculate that the rimosaviruses may be also transmitted by aphids, but given the extreme sequence differences of many of the rimosavirus RTDs from those of L/P/Es, we wonder if some rimosaviruses may be transmitted by other insect species or possibly non-insect vectors. For example, the unrelated beet necrotic yellow vein virus (*Benyviridae*) encodes an RTD extension on the CP that facilitates fungal transmission of the virus (Tamada et al., 1996). In the poleroviruses, the C-terminal half of the RTD also has been shown to play a role in virus movement in the phloem (Peter et al., 2009;Boissinot et al., 2014;Xu et al., 2018a). Thus, it appears that ORFs 3a, 4, and C-terminus of ORF5 may act together to ensure efficient, phloem-limited virus movement in the infected plant (DeBlasio et al., 2018). Other functions are possible. Recently, a CP-RTD protein of an ilarvirus was shown to have silencing suppressor activity (Lukhovitskaya et al., 2024). The role(s), if any, the rimosavirus RTD plays in vector transmission, virus movement, or silencing suppression is one of the many interesting questions about this cryptic genus that remains to be answered.

Finally, based on the phylogenetic tree showing that all L/P/E RTDs fall into one clade that is less diverse than the branches that include only rimosaviruses, we propose that rimosavirus RTDs have a very ancient origin, and/or have been undergoing more rapid selection and evolution than the L/P/E RTDs. The polero- and enamoviruses have a replication apparatus so different from the rimosaviruses and luteoviruses that they fall into a different phylum (*Pisuviricota*) from the *Tombusviridae* (*Kitrinoviricota*) (Miller and Lozier, 2022). Thus, we speculate that a sobemo-like ancestor of the polero- and enamoviruses may have acquired its RTD by recombination in mixed infection with an ancestral rimosa-like or luteo-like virus.

### 4.3 Noncanonical translation

#### 4.3.1 Potential IRES in the 5’ UTR

Perhaps the most unusual feature of rimosaviruses is the long tract at the 5’ end upstream of the ORF1 initiation codon. MaTV encodes a significant size ORF (ORF0) in this region, and PVLaTV1 encodes a truncated version of this ORF, however (i) there are AUGs upstream of ORF0, (ii) no ORF of substantial size is present or conserved upstream of ORF1 in the other rimosaviruses, and (iii) there are numerous AUGs scattered at different positions upstream of ORF1 in all rimosavirus genomes (Supplementary Fig. S1). Thus, we speculate that the 5’UTR may have IRES or ribosome shunting activity, which would be novel, because genomes in the other tombusvirid genera contain a short 5’ UTR (maximum 142 nt in BYDV), relying on the 3’ CITE that facilitates translation by ribosome scanning from the 5’ end. The rimosavirus 5’ UTR has potential to contain an IRES or shunting structure, but we found no significant, conserved secondary structures in the 5’ UTRs. A conserved G-poor tract immediately upstream of the ORF1 start codon is a feature in some translation enhancers such as the tobacco mosaic virus 5’ leader (Sleat et al., 1987), and we also found the tract upstream of and including the G-poor tract is C-rich, which is a feature in some IRESes (Pilipenko et al., 1992). Although the 3’ UTR is long enough to encode a 3’ CITE (Simon and Miller, 2013), we found no secondary structures in the 3’ UTR that resembled known CITEs. Additional computational approaches and of course lab experiments are necessary to determine how translation initiates on rimosavirus genomes.

#### 4.3.2 Leaky start and stop codons

One translation feature that is not mysterious is the secondary structure that controls readthrough of the ORF1 stop codon to allow translation of the RdRp encoded in ORF2. All twelve viruses have obvious bulged stem-loops adjacent to the ORF1 stop codon that can base pair to a 3’ distal readthrough element (DRTE) as has been shown to facilitate readthrough in many tombusviruses (Cimino et al., 2011). Also, the stem-loop near the 3’ end, with the GGGC bulge capable of pseudoknot base pairing to the extreme 3’ terminal bases GCCC resembles the replication structure present in all other studied tombusvirid genomes (Koev et al., 2002;Simon, 2015).

By analogy with luteo- and poleroviruses (Tacke et al., 1990;Dinesh-Kumar and Miller, 1993), we speculate that ORFs 4, 3 and 5 are translated from sgRNA1, initiating upstream of the ORF3 start codon, as the CP of all tombusvirids is translated from a sgRNA. If our hypothesis of non-AUG translation initiation of ORF4 is correct, then the 5’ end of sgRNA1 must be downstream of any AUG triplets located upstream of the ORF3 AUG, because if the 5’ end of sgRNA1 included those AUGs, scanning ribosomes would initiate at those, rather than the predicted non-AUG start codon and subsequent AUG initiator of ORF3. Bearing this in mind, we sought conserved sequences and secondary structures that may be required for generating the 5’ end of sgRNA1, because such structures have been shown to be required for sgRNA1 synthesis in many tombusvirids (Lin and White, 2004;Jiwan and White, 2011). Indeed, a conserved sequence and stem-loop is predicted in this region of the rimosavirus genomes (underlined bases in Supplementary Fig. S5). Co-variations in the few base differences among these sequences support the existence of the stem-loop, implying that base pairing is required, even if the sequences that comprise those base pairs vary (compare underlined regions in PVLaTV1 and AVE in Supplementary Fig. S5).

In conclusion, phylogenetic comparisons reveal that the twelve genome sequences found in GenBank described here clearly belong to viruses in a new genus in the *Tombusviridae* family. The biological properties of the viruses associated with these genomes, such as host range, symptomatology, vector specificity, remain to be elucidated. It is clear that the diverse catalog of noncanonical translation mechanisms that are a hallmark of tombusvirid gene expression is enriched by this puzzling new collection of viral genomes.

## Acknowledgments

The authors thank Megan Harrison, Seema Raychaudhuri, and Abigail Maue for assistance in preparing this manuscript.

## Supplementary Figures

**Supplementary Fig. S1**. 5’ UTR alignments using default parameters for MUSCLE version 3.8.1551 in SnapGene. Sequences end at the start codon for ORF1. Bases with ≥50% sequence identity at each position are shaded in yellow, with intensity of shading proportional to number of sequences in which the base is conserved at each position. All AUG triplets are highlighted in green.

**Supplementary Fig. S2.** Alignment of amino acid sequences of RdRp domains encoded by ORF2. Alignment was made using MUSCLE version 3.8.1551 in SnapGene using default parameters.

**Supplementary Fig. S3.** Predicted (MFOLD) secondary structures of the 3’ termini of GenBank sequences of the indicated viral genomes. HubTLV2 sequence is likely incomplete, as it does not terminate in GCCC. Predicted 3’ terminus of ENaTV10 genome is indicated. Bases we speculate are adapter-derived are in gray. In this portion of the genome, TTTV1 differs from ENaTV10 at only one base position (indicated) and the TTTV1 GenBank sequence terminates before the actual predicted 3’ end, which is the same as that predicted for ENaTV10. Dashed line indicates pseudoknot base pairing conserved in all tombusvirids (dark red and dark blue). Extended potential base pairing beyond conserved 4 base pairs is indicated in lighter shades of red and blue. Downstream translational readthrough element (DTRE), predicted to base pair to a bulge in the stem-loop that facilitates readthrough of the ORF1 stop codon are highlighted in yellow.

**Supplementary Fig. S4**. Predicted secondary structures of regions in the 5’ end of the 3’ UTRs of the indicated viruses. Sequences begin with the ORF5 stop codon.

**Supplementary Fig. S5.** Alignment of nucleotide sequences of the AUGs that might be predicted to start ORF4, except for frame changes caused by indels before ORF3 in most cases, through the start codon of ORF3, which overlaps ORF4 in a different frame. Shade of yellow highlighting increases with increased sequence identity at each position. The two underlined tracts are predicted (MFOLD) to base pair to each other to form a stem-loop that may contribute to generating the 5’ end of sgRNA1.

**Supplementary Fig. S6.** Predicted base pairing 3’ proximal to the CP ORF stop codon (UAG in red). 203 nt were used in each prediction. Base numbering starts with the first base of the stop codon. Only the UAG-proximal helices are shown. Dashed line indicates long structured region not shown. TTTV1 and ENAV10 sequences are identical with the exception of the blue U in ENaTV10 vs A in TTTV1 at position 6.

