## SupplementalAlignmentsandStructures for "A proposed new *Tombusviridae* genus featuring extremely long 5’ untranslated regions and a luteo/polerovirus-like gene block"

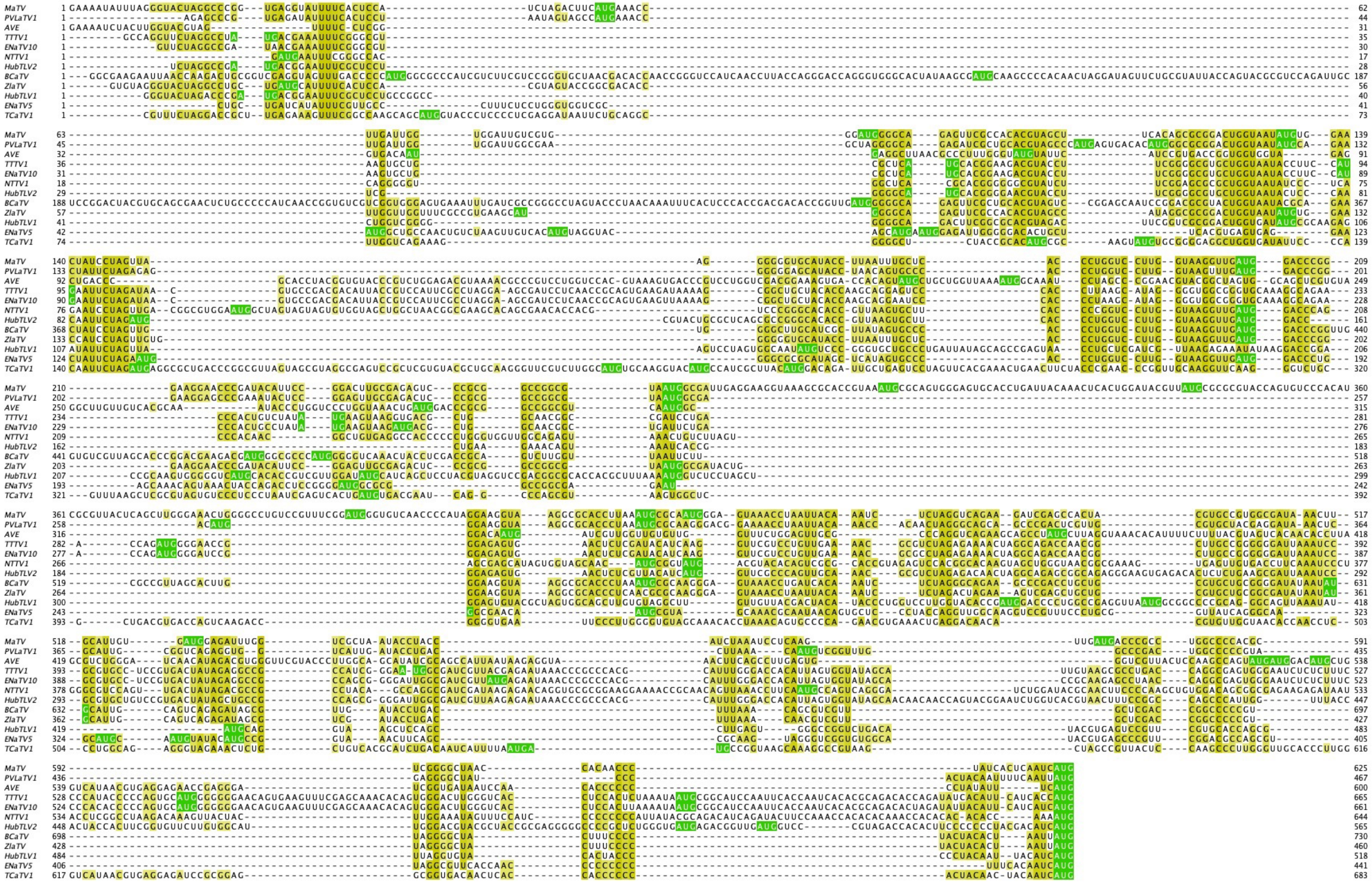

Supplementary Fig. S1

Alignments of 5' UTRs

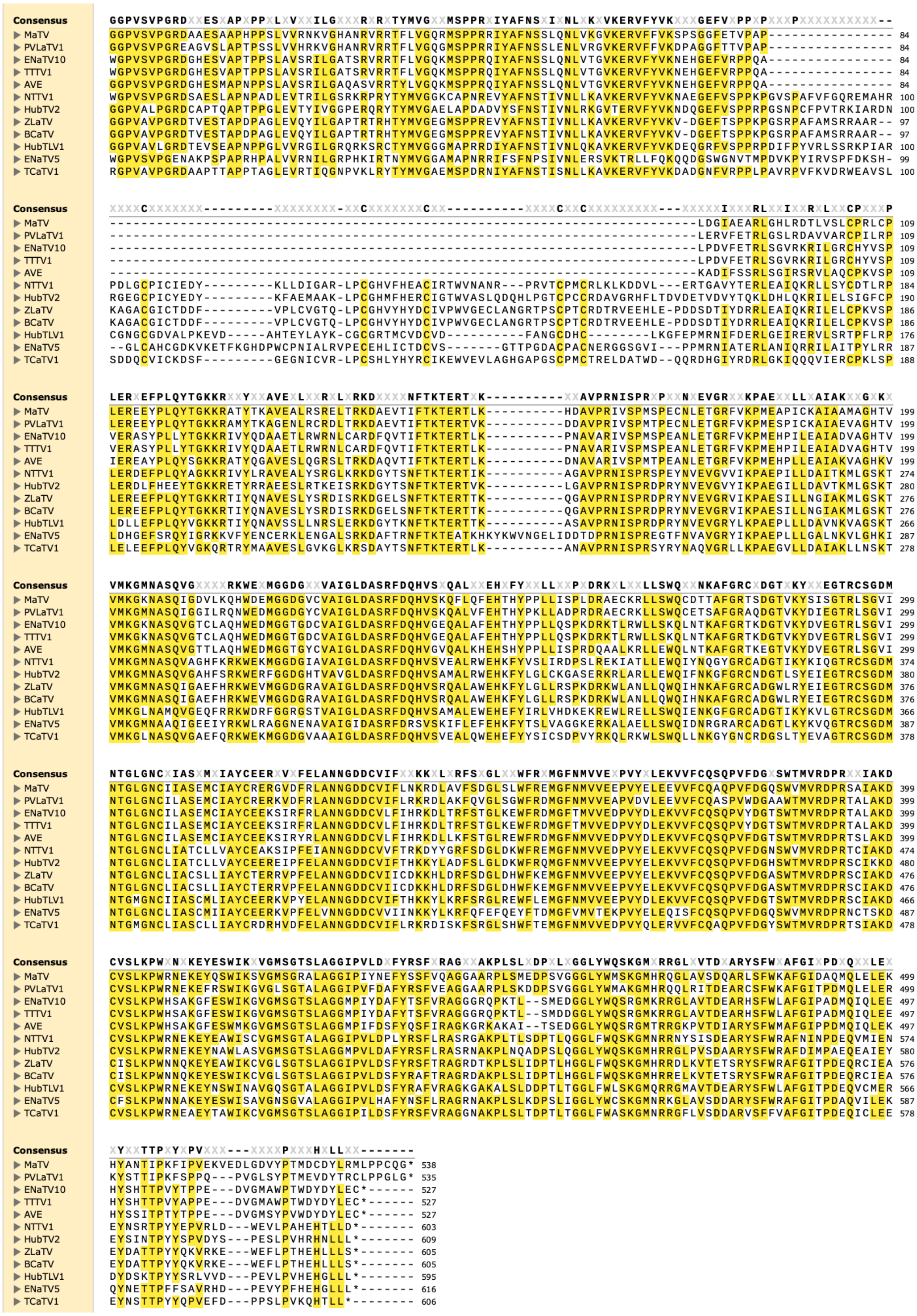

Supplementary Fig. S2. Alignment of RdRp domains encoded by ORF2

Predicted 3'-terminal secondary structures

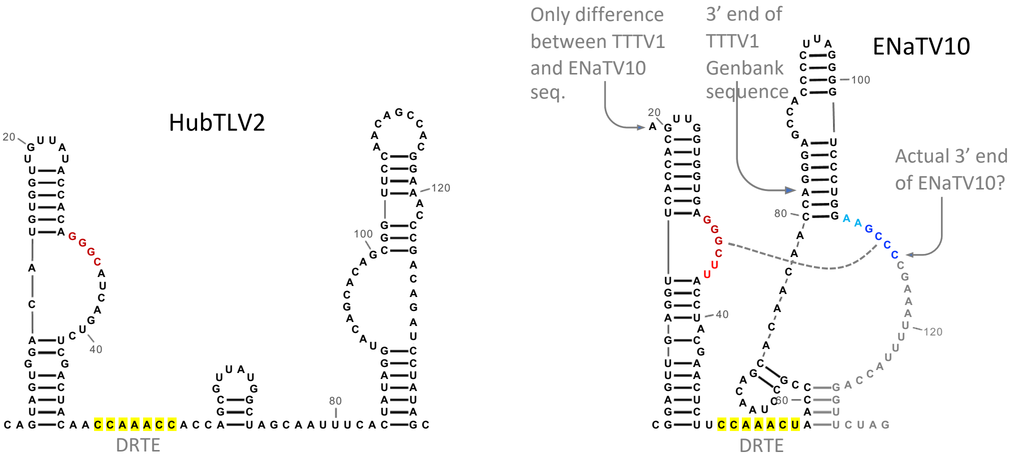


Supplemental Fig. S3

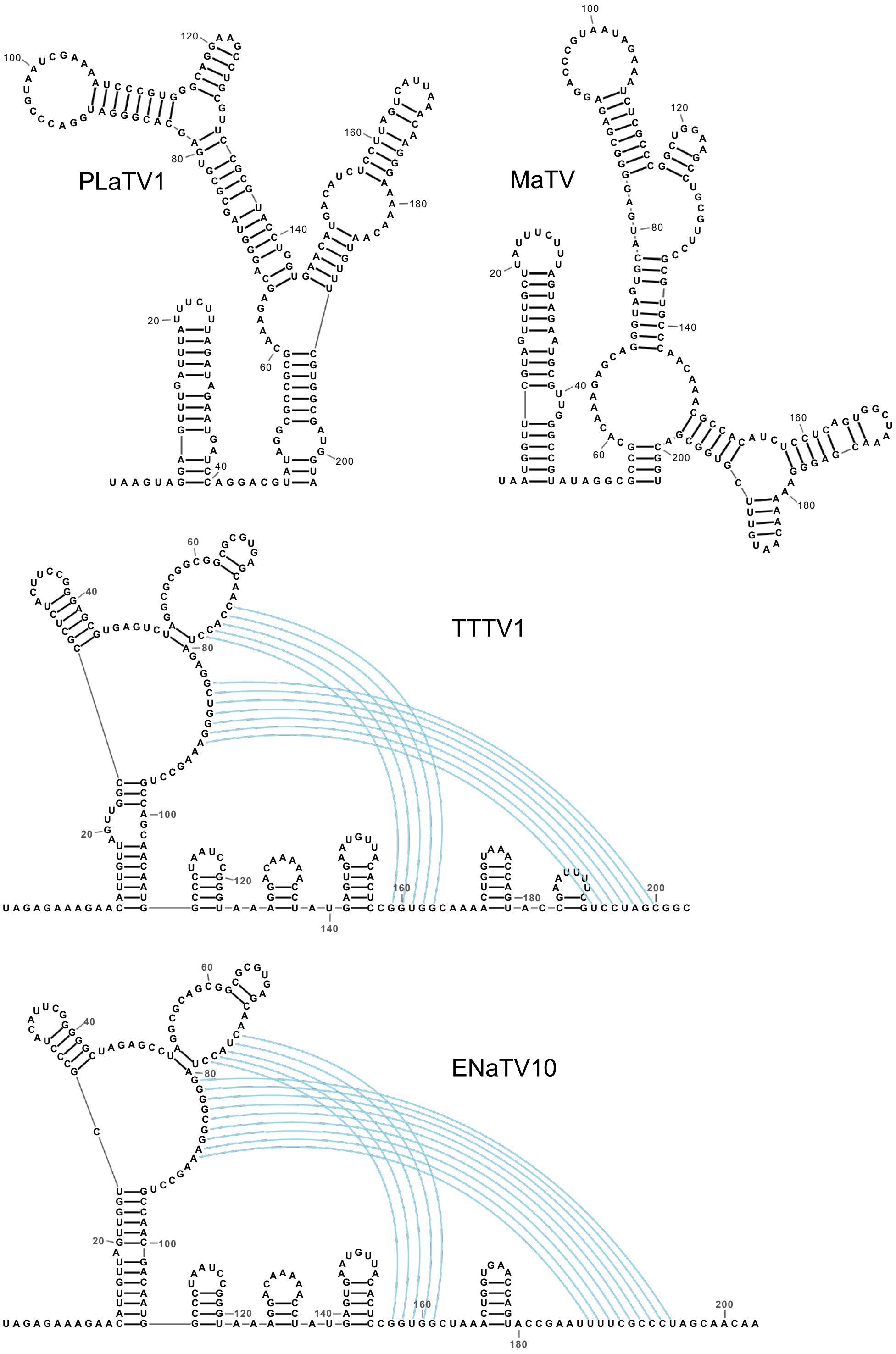


Supplementary Fig. S4

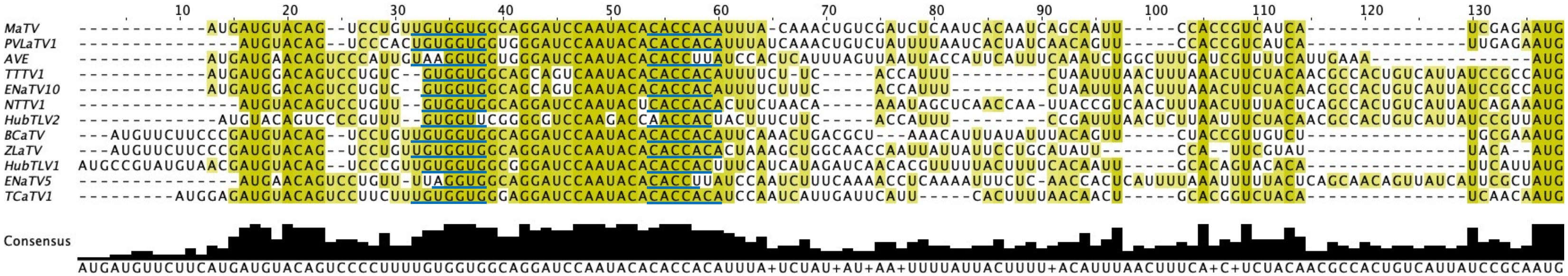


Supplementary Fig. S5

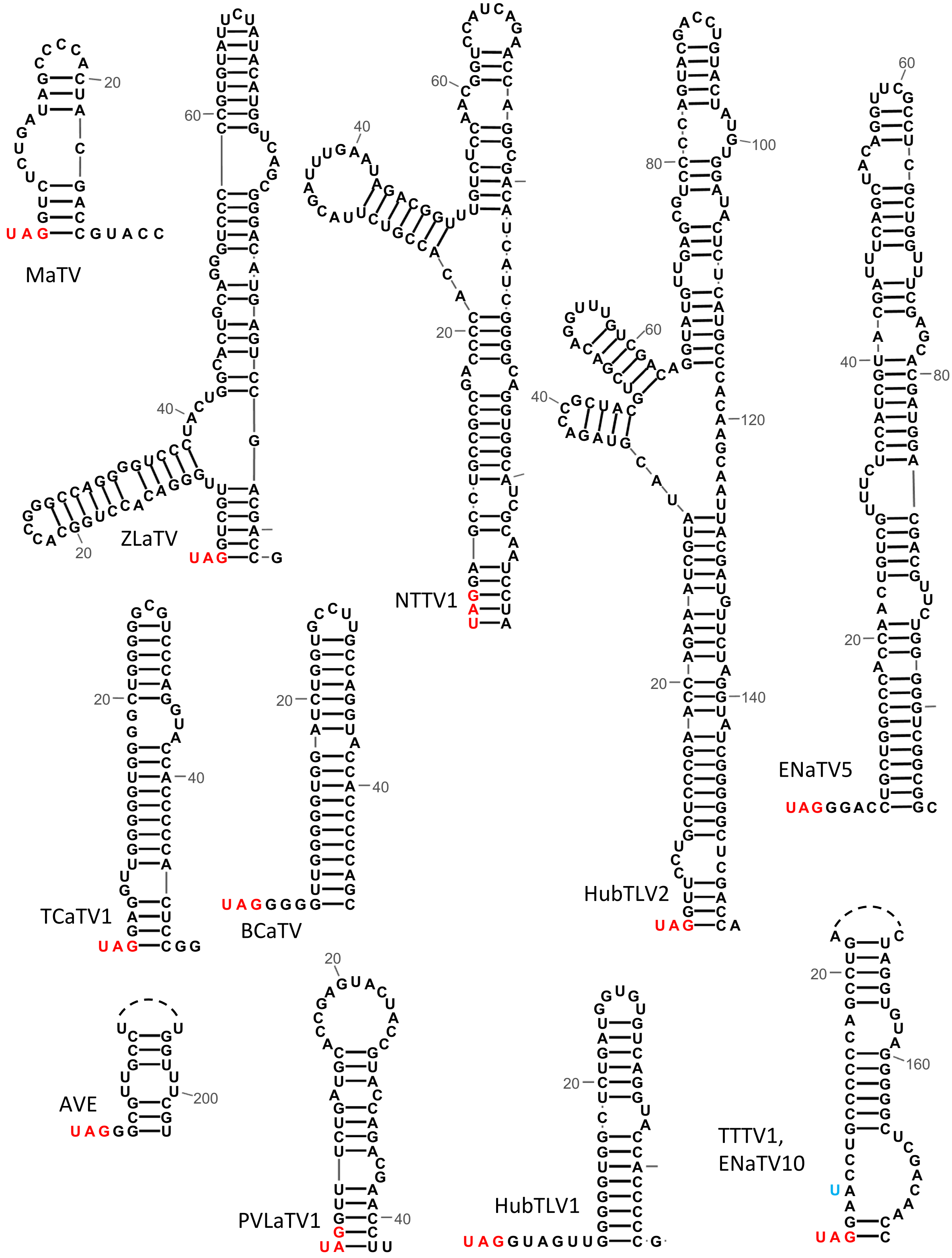


Supplementary Fig. S6

**Supplementary Table S1**

<u>Custom 5′ RACE oligodeoxynucleotides</u>
Nested primer 1: `AACGCGATGTGGGACACTGGTACGCGC`
Nested primer 2: `ATCGCCATTACGCCGGCCGCGGGACT`
Nested primer 3: `GGCCGCGGGACTCTCGCAAGTCCGGAATGTATCGGG`
5R1: `TCGCAAGTCCGGAATGTAT`

<u>Custom 3′ RACE oligodeoxynucleotides</u>
3′poly(A): `AAAAAAAAAAAAAAAAGTCGACATCG`
3F1: `CGTATAGGCGCCGCACAAAGAG`
3F2: `TAATAGAAATCTCGCCCGGCTGGA`

**Supplementary Table S2.** Molecular weights (kDa) of proteins encoded by the indicated open reading frames (ORFs).

| Viral Genome | ORF0 | ORF1 | ORF2 | ORF3 | ORF4[a] | ORF5 |
|---|---|---|---|---|---|---|
| MaTV | 11.6 | 25.7 | 59.7 | 22.9 | ~19 | 39.6 |
| PVLaTV1 | 6.2 | 26.1 | 59.3 | 23.5 | ~19 | 38.9 |
| AVE | NA | 30.7 | 58.7 | 23.6 | ~20 | 34.5 |
| TTTV1 | NA | 23.3 | 59 | 25.1 | ~17 | 37.4 |
| ENaTV10 | NA | 23.2 | 59 | 25.2 | ~17 | 37.3 |
| NTTV1 | NA | 24.8 | 68.2 | 25.8 | ~19 | 34.1 |
| HubTLV2 | NA | 25.7 | 68.7 | 24.1 | ~17 | 37.4 |
| BCaTV | NA | 25.6 | 68.2 | 22.6 | ~18 | 39.6 |
| ZLaTV | NA | 25.6 | 68.3 | 23.6 | ~20 | 40.9 |
| HubTLV1 | NA | 25.9 | 67.3 | 22.7 | ~18 | 39.6 |
| ENaTV5 | NA | 34.1 | 69.2 | 24.2 | ~18 | 34.7 |
| TCaTV1 | NA | 23.8 | 68.0 | 22.2 | ~18 | 39.2 |

[a]Estimated molecular weight assuming initiation at a known non-AUG start codon shortly upstream of ORF3 start codon.
